# Reward integration in prefrontal-cortical and ventral-hippocampal nucleus accumbens inputs cooperatively modulates engagement

**DOI:** 10.1101/2024.06.27.601063

**Authors:** Eshaan S. Iyer, Peter Vitaro, Serena Wu, Jessie Muir, Yiu Chung Tse, Vedrana Cvetkovska, Rosemary C. Bagot

## Abstract

The NAc, a highly integrative brain region controlling motivated behavior, is thought to receive distinct information from various glutamatergic inputs yet strong evidence of functional specialization of inputs is lacking. While circuit neuroscience commonly seeks specific functions for specific circuits, redundancy can be highly adaptive and is a critical motif in circuit organization. Using dual-site fiber photometry in an operant reward task, we simultaneously recorded from two NAc glutamatergic afferents to assess circuit specialization. We identify a common neural motif that integrates reward history in medial prefrontal cortex (mPFC) and ventral hippocampus (vHip) inputs to NAc. Then, by systematically degrading task complexity, dissociating reward from choice and action, we identify key circuit-specificity in the behavioral conditions that recruit encoding. While mPFC-NAc invariantly encodes reward, vHip-NAc encoding is uniquely anchored to loss. Ultimately, using optogenetic stimulation we demonstrate that both inputs co-operatively modulate task engagement. We illustrate how similar encoding, with differential gating by behavioral state, supports state-sensitive tuning of reward-motivated behavior.

## INTRODUCTION

Redundancy is a defining property of nervous system organization (Marder, 2011; Marder & Goaillard, 2006; Mizusaki & O’Donnell, 2021) yet there has been limited consideration of the role of redundancy in neural circuit mechanisms of motivated behavior. Redundancy in neural circuits may confer various advantages, including increasing robustness of cognition and behavior to perturbation, enhancing encoding accuracy, and facilitating coherent integration of multiple inputs, suggesting it should be a frequently observed motif (Ghanbari et al., 2023; Hiratani & Fukai, 2018; Nguyen et al., 2019). While the literature abounds with examples of apparent circuit-specific cognitive and behavioral functions, the potential for redundancy is rarely examined. To better understand these opposing motifs in nucleus accumbens (NAc) circuits we leveraged dual circuit recordings and computational modeling to rigorously test the specificity and redundancy of information processing in a fully controlled, within animal comparison of two NAc glutamatergic inputs.

The NAc integrates glutamatergic inputs with dopaminergic input from the ventral tegmental area, with multiple glutamatergic inputs converging at the level of individual medium spiny neurons in the NAc medial shell (Britt et al., 2012; Carter et al., 2007; Christoffel et al., 2021; Floresco, 2015; French & Totterdell, 2002; Lind et al., 2023; Muir et al., 2024; O’Donnell & Grace, 1995). Prominent theoretical perspectives hold that these inputs send qualitatively distinct information which the NAc then integrates to orchestrate motivated behavior (Floresco, 2015; Grace et al., 2007; Lind et al., 2023; Parker et al., 2022). For example, the mPFC contributes information about rewarding events and executive control while the vHip contributes emotional context and behavioral inhibition (Bagot et al., 2015; Barker et al., 2019; Hamel et al., 2022; Lindenbach et al., 2022; Muir et al., 2020; Otis et al., 2017; Parker et al., 2022; Spellman et al., 2021; Wenzel et al., 2023; Yoshida et al., 2020). Despite predictions of distinct encoding and behavioral function for mPFC-NAc and vHip-NAc, strong evidence of functional specialization is lacking. To date, most studies have examined a single input and the few studies that examined one or more inputs in the same task compared across animals leaving open the possibility that inter-individual variation in behavior and other variables influence neural encoding (Britt et al., 2012; Chen et al., 2023; Reed et al., 2018).

To systematically interrogate functional redundancy versus specialization we simultaneously probed neural encoding using dual-site *in vivo* fiber photometry to record activity in two glutamatergic circuits during reward-guided choice in a two-armed bandit task. The mPFC-NAc is widely appreciated to mediate reward processing and, given that vHip-NAc inputs converge with mPFC-NAc, we asked if the vHip-NAc might also contribute to this function (Otis et al., 2017; Parker et al., 2022; Spellman et al., 2021). Using trial-by-trial modeling of neural activity, we identify a novel mechanism for integrating outcome information across trials that is common to both circuits. Analyzing the redundancy across signals revealed an additional dimension of uniqueness to vHip-NAc encoding. By sequentially degrading task complexity we show that, despite sharing a common mechanism for outcome integration, each circuit is recruited in distinct behavioral states, with the vHip-NAc preferentially encoding reward after loss.

Optogenetically manipulating circuit-specific activity revealed that, once recruited, both inputs cumulatively mediate dynamic behavioral engagement. Our findings reveal co-operative circuit organization in NAc wherein redundant encoding in two inputs is gated by circuit-specific mechanisms for state-sensitive tuning of reward-motivated behavior.

## Results

### mPFC-NAc and vHip-NAc similarly encode outcomes in a probabilistically rewarded environment

To assess redundancy versus specificity in outcome encoding in two distinct circuits under matched conditions and trial histories, we injected retrograding AAV-GCaMP7f in NAc medial shell and implanted optic fibers in mPFC and vHip to record Ca^2+^-associated fluorescence while mice engaged in reward-guided choice (Fig. 1E). We trained mice in a two-lever probabilistic reward learning task (i.e. a two-armed bandit task) in which lever pressing probabilistically earns a chocolate milk reward (Fig. 1A). Following each lever press, one of two different auditory cues signaled trial outcome (rewarded, unrewarded) and start of the inter-trial interval (ITI). To maintain a dynamic environment with robustly encountered rewarded and unrewarded outcomes, levers were probabilistically rewarded on 80% or 20% of presses with probabilities switched after five consecutive responses on the high probability lever. Female (n=10) and male (n=12) mice experienced similarly high numbers of unrewarded and rewarded trials and low numbers of omission trials (Fig. 1B-D). Examining behavior across sessions shows decreasing staying probability after unrewarded outcomes and increasing rewards earned, indicating animals use information about outcomes to guide behavior (Fig. S1).

**Figure 1.**
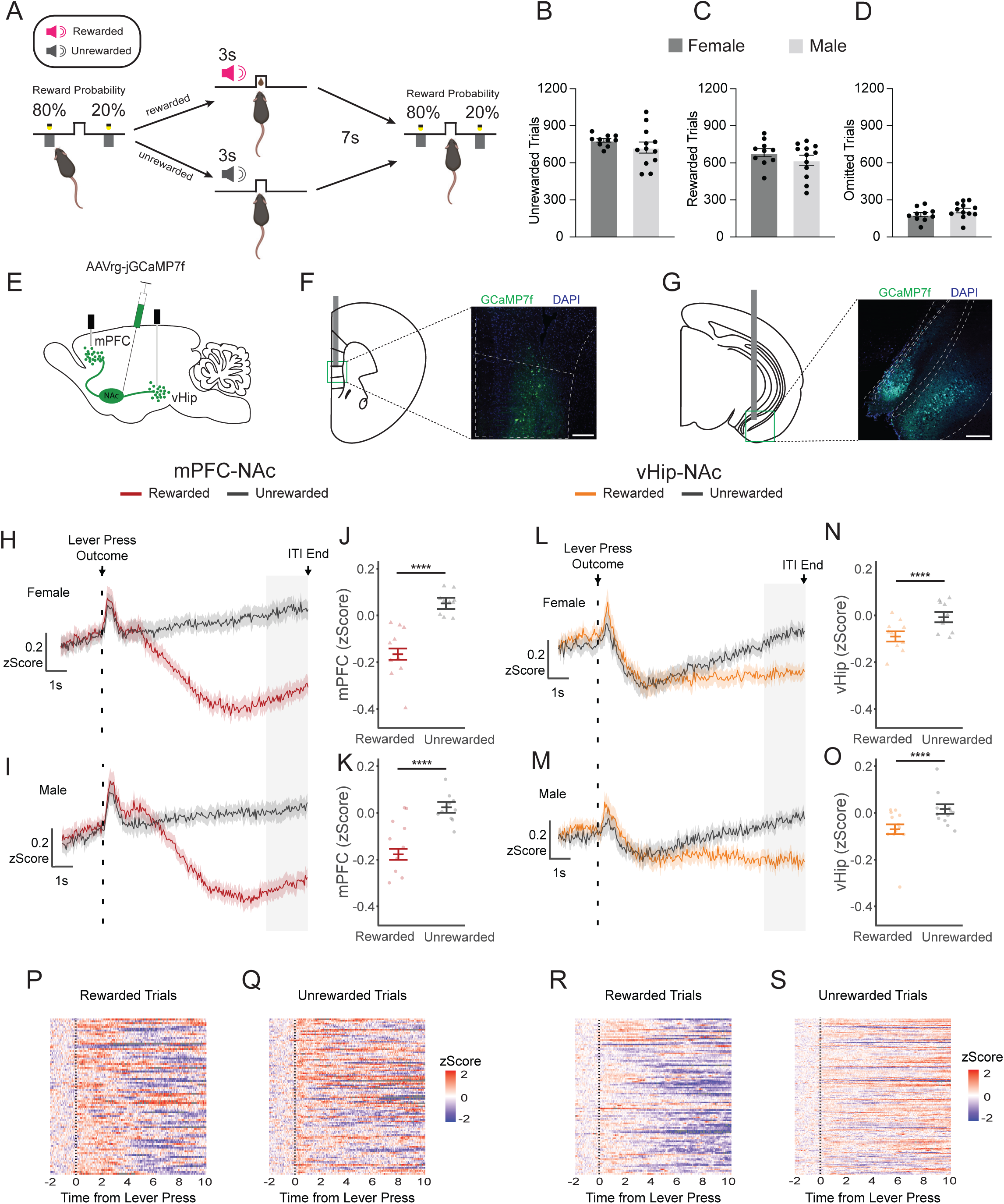
mPFC-NAc and vHip-NAc similarly encode reward in a probabilistically rewarded environment. (A) Schematic of two-armed bandit task. Mice lever press in a two lever task in which one lever is rewarded with chocolate milk on 80% of trials, and the other on 20%. Following a lever press, levers retract, and auditory cues signal outcome and start of a 10 sec inter-trial interval (ITI). Contingencies switch after five consecutive responses on the high probability lever. Female (n=10) and male (n=12) mice robustly engage with the task, experiencing similar numbers of (B) unrewarded (C) rewarded and (D) omission trials. (E) Retrograding jGCaMP7f is injected into the nucleus accumbens (NAc) and optic fibers implanted in medial prefrontal cortex (mPFC) and ventral hippocampus (vHip) to simultaneously probe neural activity indicated by Ca^2+^-associated fluorescence changes in (F) mPFC neurons projecting to NAc (mPFC-NAc) and (G) vHip neurons projecting to NAc (vHip-NAc) as mice encounter reward and non-reward. Estimated mean mPFC-NAc activity across all rewarded and unrewarded trials in (H) female (n=10) and (I) male (n=12) mice. Analysis focused on 8-10 sec after lever press (ITI end). At ITI end, mPFC-NAc activity is suppressed by rewarded outcomes in female (J; Z=21.348, p<0.0001) and male (K; Z=19.625, p<0.0001) mice. Estimated mean vHip-NAc activity across all rewarded and unrewarded trials in (L) female and (M) male mice. At ITI end, vHip-NAc activity is suppressed by rewarded outcomes in female (N; Z=8.161; p<0.0001) and male (O; Z=8.924; p<0.0001) mice. Heatmap of mPFC-NAc activity to (P) rewarded outcomes and (Q) unrewarded outcomes in a representative animal across one session. Heatmap of vHip-NAc activity to (R) rewarded outcomes and (S) unrewarded outcomes in a representative animal across one session. Error bars represent SEM around the estimated mean. ****p<0.0001

Trial based tasks are ideal for probing neural encoding, generating large numbers of trials. However, standard analysis approaches either analyze individual trials, failing to account for the within animal nested data structure and inappropriately inflating effects, or average all trials within animals, thereby underestimating effects. Choice tasks are additionally challenging with the number of instances of each trial type varying across animals. To preserve the power of trial-by-trial data while accounting for the nested structure and unbalanced observations we used a linear mixed model approach (Yu et al., 2022). To examine how outcome is encoded in each projection, we modeled normalized Ca^2+^-associated fluorescence change as a function of trial outcome, while controlling for inter-individual variability.

Reward strongly suppressed mPFC-NAc and vHip-NAc activity in female and male mice. In mPFC-NAc, a peak following the lever press and outcome delivery is followed by gradually emerging reward-associated suppression across the ITI (Fig 1H,J,P,Q). In vHip-NAc, an initial peak is followed by suppression after the lever press and outcome delivery, with suppression sustained following reward or activity gradually increasing following unrewarded outcomes (Fig. 1 L,N,R,S). We focused analysis on the end of the ITI (8-10 sec after lever press) when trial outcome has been integrated prior to next trial start. By ITI end, reward robustly suppressed mPFC-NAc activity in female and male mice (Fig. 1I,K). Reward also robustly suppressed vHip-NAc activity in female and male mice (Fig. 1M,O). This indicates that outcome encoding emerges across the ITI with reward suppressing mPFC-NAc and vHip-NAc activity. To explore modulation by other task factors, we examined neural encoding time-locked to licking and decision-relevant behaviors. We did not observe clear neural encoding of licking (Fig. S2A,D), the identity of the chosen lever (Fig. S2B,E), or the decision to stay or shift (Fig. S2C,F) suggesting that outcome is the primary source of modulation in mPFC-NAc and vHip-NAc in this task.

Having observed that mPFC-NAc and vHip-NAc are similarly modulated by reward, we then examined if one circuit leads the other. Calculating the time lag for the maximum cross-correlation, we found that the regression coefficients of the mPFC-NAc and vHip-NAc cross-correlation did not significantly differ from zero in rewarded or unrewarded trials in either sex. This shows that neither circuit drives outcome encoding in the other (Fig. S3A, B). However, we observe that suppression emerges earlier in vHip-NAc than mPFC-NAc, suggesting that while overall encoding might be similar, the underlying dynamics likely vary between the pathways (Fig. S3C).

### mPFC-NAc and vHip-NAc similarly integrate reward history

We find that mPFC-NAc and vHip-NAc similarly encode outcomes. Observing that reward-induced suppression lasted throughout the ITI, we speculated that this enduring modulation might integrate reward information across successive trials and that this integration might be more prominent in mPFC-NAc than vHip-NAc, given prior evidence of enduring representation in mPFC (Parker et al., 2022; Spellman et al., 2021; Sul et al., 2010). To test this, we sorted trials by both prior and current outcome, identifying trial sequences that were rewarded then rewarded (R→R), rewarded then unrewarded (R→U), unrewarded then rewarded (U→R), and unrewarded then unrewarded (U→U). We then compared neural activity across the ITI on the most recent trial to determine how prior outcome modulates outcome encoding on the current trial. Analyzing males and females separately revealed similar modulation (Fig. S4) and we therefore report sex-combined analyses. Both previous and current outcome modulate mPFC-NAc activity (Fig. 2A). Following a given trial (t-1), reward suppresses mPFC-NAc activity (Fig. 2B) effectively resetting the baseline for the next trial. Reward on the subsequent trial (t0) similarly suppresses mPFC-NAc by ITI end, regardless of prior outcome. However, when mice are unrewarded on the subsequent trial (t0), suppression of mPFC-NAc by prior reward is maintained through ITI end (Fig. 2C). This suggests that a single reward maximally and enduringly suppresses mPFC-NAc activity and that, in the absence of subsequent reward, this suppression slowly dissipates.

**Figure 2.**
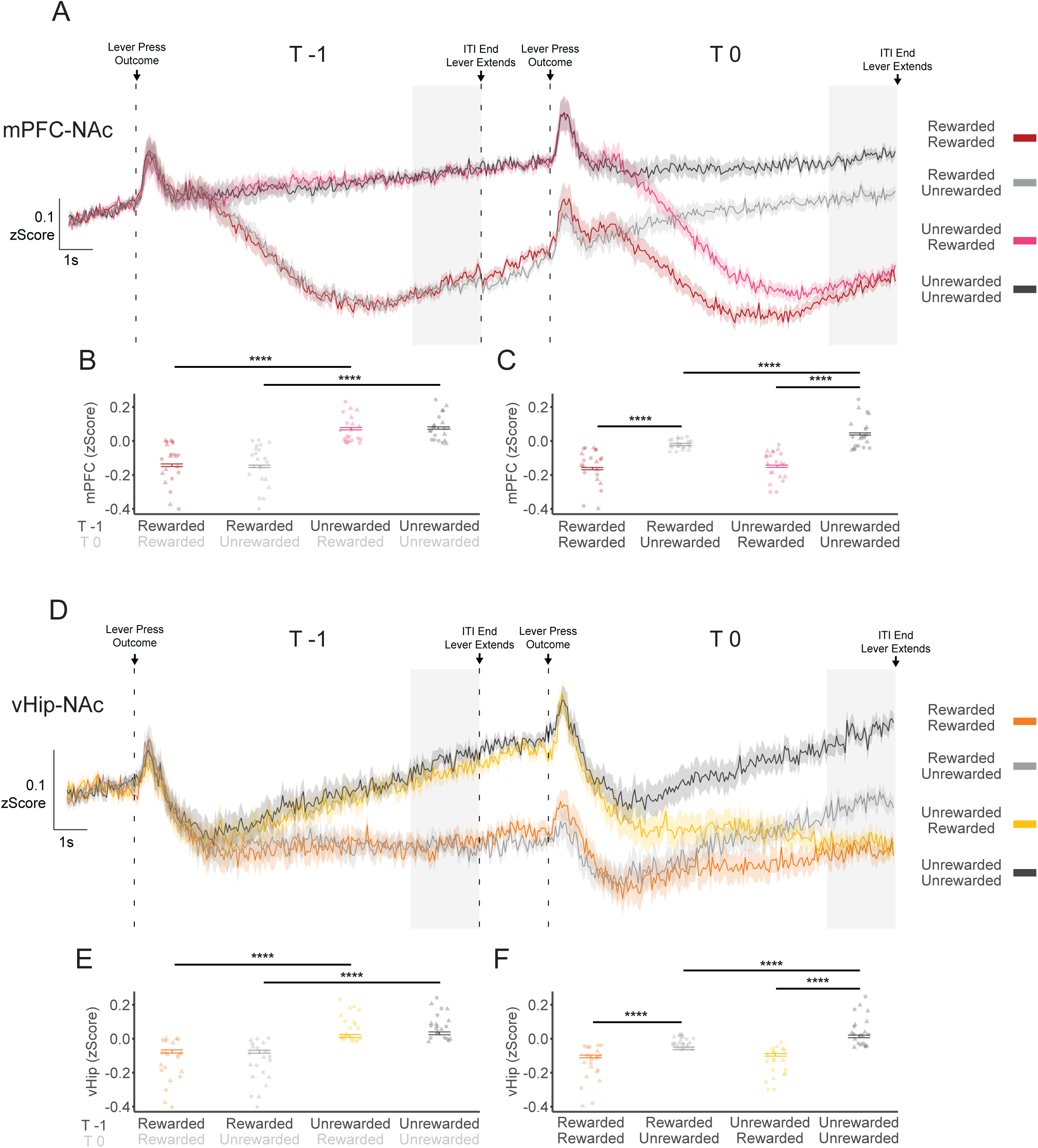
mPFC-NAc and vHip-NAc similarly integrate reward history. (A) Estimated mean mPFC-NAc activity across pairs of consecutive trials (t-1→t0) showing rewarded+rewarded (R→R), rewarded+unrewarded (R→U), unrewarded+rewarded (U→R), and unrewarded+unrewarded (U→U) trial pairs in female (n=10) and male (n=12) mice. Analysis focused on 8-10 sec after lever press (ITI end). (B) On trial t-1, mPFC-NAc activity is significantly suppressed by reward (U→U vs R→U: Z=28.99496, p<0.0001; U→R vs R→R: Z=25.6767, p<0.0001). (C) On the subsequent trial, t0, mPFC-NAc activity is significantly suppressed by current reward (U→U vs U→R: Z=-28.5098, p<0.0001; R→U vs R→R: Z=-19.8981, p<0.0001). When trial t0 is unrewarded, mPFC-NAc activity remains significantly suppressed by reward experienced on the previous trial, t-1, (U→U vs R→U: Z= 9.19653, p<0.0001). (D) Estimated mean vHip-NAc activity across pairs of consecutive trials (t-1→t0) showing rewarded+rewarded (R→R), rewarded+unrewarded (R→U), unrewarded+rewarded (U→R), and unrewarded+unrewarded (U→U) trial pairs. (E) On trial t-1, vHip-NAc activity is significantly suppressed by reward (U→U vs R→U: Z=14.9372, p<0.0001; U→R vs R→R: Z=11.6962, p<0.0001). (F) On the subsequent trial, t0, vHip-NAc activity is significantly suppressed by current reward (U→U vs U→R: Z=-17.4993, p<0.0001; U→U vs R→R: Z= −7.1005, p<0.0001). When trial t0 is unrewarded, vHip-NAc activity remains suppressed by reward experienced on the previous trial, t-1, (U→U vs R→U: Z= 11.4112, p<0.0001). Individual-animal averages are indicated by circles for males and triangles for females. Error bars represent SEM around the estimated mean. ****p<0.0001

We then examined if vHip-NAc similarly integrates outcomes (Fig. 2D). Following a given trial (t-1), reward suppresses vHip-NAc activity (Fig. 2E). As with mPFC-NAc, this resets the baseline for the next trial (t0), wherein reward suppresses vHip-NAc regardless of prior outcome.

However, when the subsequent trial (t0) is unrewarded, suppression of vHip-NAc activity by prior reward is maintained through ITI end (Fig. 2F). Together, this shows that mPFC-NAc and vHip-NAc similarly integrate outcomes across trials. In both circuits, reward maximally suppresses neural activity and activity gradually increases following subsequent unrewarded outcomes, such that, by ITI end, the relative degree of suppression represents an integrated reward outcome history.

### mPFC-NAc and vHip-NAc are differentially sensitive to loss

Analyzing neural encoding of reward and outcome integration revealed that mPFC-NAc and vHip-NAc similarly encode reward suggesting they may provide redundant information to the NAc. To test redundancy between mPFC-NAc and vHip-NAc we calculated the conditional entropy of mPFC-NAc given vHip-NAc (□(mPFC-NAc|vHip-NAc)) and vHip-NAc given mPFC-NAc (□(vHip-NAc|mPFC-NAc)). In this way, we assessed the information contributed by each circuit beyond that contributed by the other at ITI end, when outcome is fully integrated (Fig. 3A). We contrasted entropy between rewarded and unrewarded outcomes as a function of prior outcome. Relative to unrewarded outcomes, the entropy of mPFC-NAc given vHip-NAc was reduced by rewarded outcomes, indicating that vHip-NAc and mPFC-NAc signals are more redundant after reward than non-reward (Fig. 3B). In contrast, following previous unreward, but not previous reward, current reward increased the entropy of vHip-NAc given mPFC-NAc (Fig. 3C), indicating that, under these conditions, mPFC-NAc explains less of the vHip-NAc signal. This shows that, after reward, vHip-NAc and mPFC-NAc encoding converges, becoming more redundant, but when reward is made more surprising by immediately following loss, vHip-NAc carries additional information. That is, despite global redundancy in reward encoding motifs, we identify a dimension of circuit specificity and a potential unique role for vHip-NAc in encoding reward following loss.

**Figure 3.**
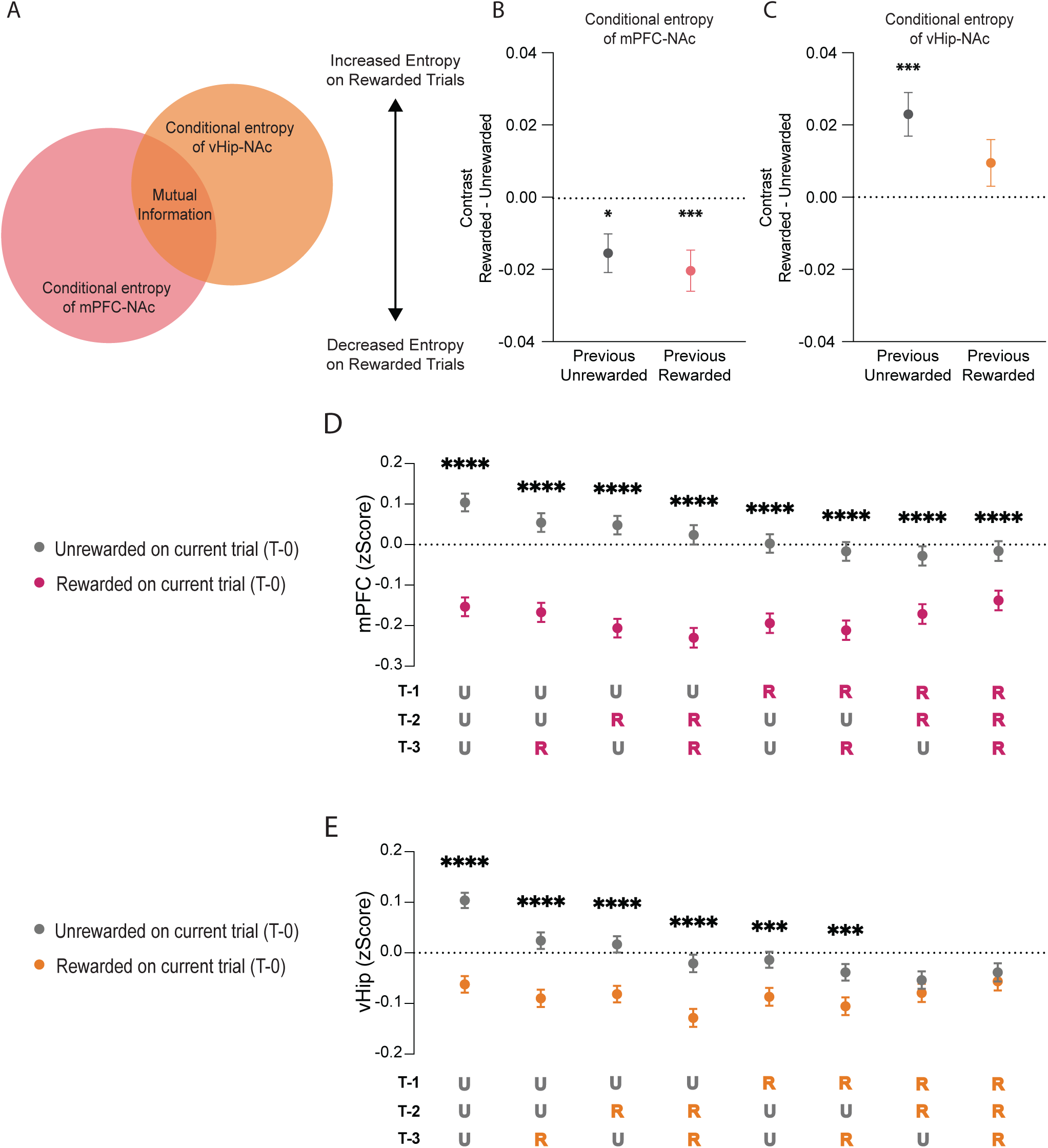
mPFC-NAc and vHip-NAc are differentially sensitive to loss. (A) Venn diagram representing the relationship between the mutual information and conditional entropy that exists between observed mPFC-NAc and vHip-NAc signals. Conditional entropy is a measure of the additional unique information contributed by a second signal given fully knowledge of a first signal. (B) Conditional entropy in mPFC-NAc is reduced on rewarded relative to unrewarded trials regardless of previous outcome (U→U vs U→R: Z=2.8644, p<0.0001; R→U vs R→R: Z=3.5185, p<0.0001) indicating that less unique information is carried in mPFC-NAc after reward. (C) Conditional entropy in vHip-NAc is increased on rewarded relative to unrewarded trials only when the prior outcome was unrewarded (U→U vs U→R: Z=-3.7566, p=0.0003) indicating that more unique information is carried in vHip-NAc when reward follows nonreward. Comparison of activity at ITI end on currently rewarded or unrewarded trials considering prior outcome history up to three trials back shows that (D) mPFC-NAc activity is suppressed on every currently rewarded trial indicating that mPFC-NAc consistently encodes current outcome via relative suppression regardless of outcome history. In contrast, (E) vHip-NAc activity is suppressed on currently rewarded trials except when current reward is preceded by two (Z=1.2310, p=0.8606) or three (Z=0.8398, p=0.9834) prior consecutive rewards indicating that vHip-NAc ceases to encode current outcome via relative suppression after consistent reward. See supplementary table 29 for all comparisons. Error bars represent SEM around the estimated mean. *p<0.05, ***p<0.001, ****p<0.0001

If this is true, across outcome histories, vHip-NAc encoding should be most apparent when reward follows an unrewarded outcome, whereas, following consecutive rewards, vHip-NAc should become insensitive to outcome as rewards become less surprising. In contrast, mPFC-NAc encoding is predicted to be relatively invariant across outcome histories. To test this, we examined current outcome encoding at ITI end while considering prior outcomes up to three trials back. Consistent with our prediction, mPFC-NAc encoded current outcome regardless of prior outcome history (Fig. 3D; Supplementary Table 1) while vHip-NAc failed to encode current outcome after two or more consecutive rewards (Fig. 3E; Supplementary Table 1). This supports our hypothesis that vHip-NAc encoding anchors to loss such that it is specifically engaged by surprising rewards whereas mPFC-NAc invariantly encodes outcome.

### Degrading task requirements reveals circuit-specific roles in reward integration

Analyzing informational redundancy and encoding across varying outcome histories suggested that, while mPFC-NAc and vHip-NAc encode and integrate reward via a common mechanism, each may nevertheless serve distinct functions in reward processing. To isolate the specific conditions under which each circuit integrates outcomes we recorded neural activity while degrading task requirements to sequentially eliminate choice and action. We first eliminated choice, extending only a single lever while maintaining the requirement to press to elicit an outcome. To hold outcome experience constant, the specific sequence of reward and unreward was yoked to each animal’s prior performance on the two-lever task (Fig. 4A). In the absence of choice, mPFC-NAc continued to encode previous and current outcome (Fig. 4B). On trial t0, by ITI end, current and prior outcomes were encoded, as in the two-lever task (Fig. 4C, Fig. 2C).

**Figure 4.**
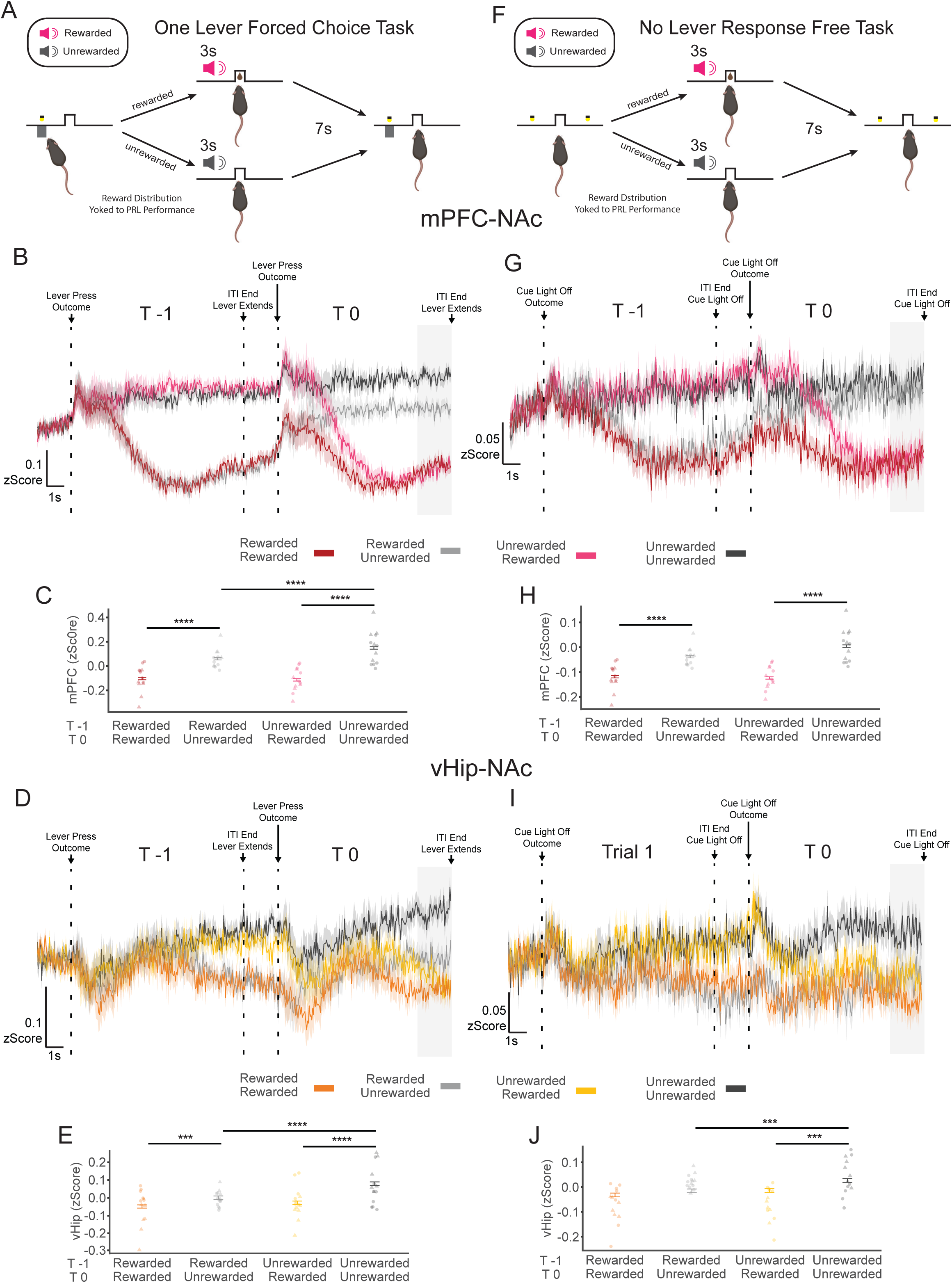
Degrading task requirements reveals circuit specialization in integrating reward history. (A) Schematic of one-lever forced choice task in which lever presses are rewarded on a schedule yoked to each animal’s individual performance in the final three days of the two-armed bandit task. Following a lever press, levers retract, and auditory cues signal outcome and start of a 10 sec inter-trial interval (ITI). (B) Estimated mean mPFC-NAc activity across pairs of consecutive trials (t-1→t0) showing rewarded+rewarded (R→R), rewarded+unrewarded (R→U), unrewarded+rewarded (U→R), and unrewarded+unrewarded (U→U) trial pairs (male n= 8, female n=6). Analysis focused on 8-10 sec after lever press (ITI end). (C) On trial t0, mPFC-NAc activity is suppressed by reward (U→U vs U→R: Z=18.8757, p<0.0001; R→U vs R→R: Z=12.0687, p<0.0001). When trial t0 is unrewarded, mPFC-NAc activity remains suppressed by reward experienced on the previous trial, t-1, (U→U vs R→U: Z= 6.3467, p<0.0001). (D) Estimated mean vHip-NAc activity across pairs of consecutive trials (t-1→t0) showing R→R, R→U, U→R, and U→U trial pairs (male n= 8, female n=6). (E) On trial t0, vHip-NAc activity is suppressed by reward (U→U vs U→R: Z=8.5245, p<0.0001; R→U vs R→R: Z=4.0519, p=0.0001). When trial t0 is unrewarded, mPFC-NAc activity remains suppressed by reward experienced on the previous trial, t-1, (U→U vs R→U: Z= 6.2425, p<0.0001). (F) Schematic of no lever response free task. Mice are allowed to collect rewards delivered on a schedule yoked to each animal’s individual trial statistics (latency and outcome) of the two-armed bandit task. Trial structure is signaled by cue-light illumination and after a predetermined delay auditory cues signal outcome and start of a 10 sec ITI. (G) Estimated mean mPFC-NAc activity across pairs of consecutive trials (t-1→t0) showing R→R, R→U, U→R, and U→U trial pairs (male n= 8, female n=6). (H) On trial t0, mPFC-NAc activity is suppressed by reward (U→U vs U→R: Z=8.2136, p<0.0001; R→U vs R→R: Z=7.4647, p<0.0001). (I) Estimated mean vHip-NAc activity across pairs of consecutive trials (t-1→t0) showing R→R, R→U, U→R, and U→U trial pairs (male n= 8, female n=6). (J) On trial t0, vHip-NAc activity is suppressed by reward only if trial t-1 was unrewarded (U→U vs U→R: Z=3.7413, p=0.0003). When trial t0 is unrewarded, vHip-NAc activity remains suppressed by reward experienced on the previous trial, t-1, (U→U vs R→U: Z= 3.8661, p=0.0002). Individual-animal averages are indicated by circles for males and triangles for females. Error bars represent SEM around the estimated mean. ***p<0.001, ****p<0.0001

Examining vHip-NAc in the one-lever task also revealed largely similar outcome-mediated modulation (Fig. 4D, Fig. 2D). At ITI end, prior and current outcomes were integrated, similar to the two-lever task (Fig. 4E, Fig. 2F). Despite conserved information encoding in both circuits, the shape of the vHip-NAc signal was more visibly altered than the mPFC-NAc. In particular, the vHip-NAc signal in the one-lever task appeared noisier and blunted with the expected peak following lever press largely absent, potentially suggesting heightened sensitivity to task structure. Overall, we find that both mPFC-NAc and vHip-NAc maintain similar graded representations of reward history that are largely independent of choice requirements.

Removing lever choice minimally impacted reward integration. We then asked if neural integration of outcome history is entirely independent of response requirements by removing both levers in a choice-free response-free task. Trials continued to be signaled by cue-lights, but without lever extension and outcomes were passively delivered yoked to each animal’s individual performance on the full two-lever task (Fig. 4F). Eliminating the response requirement markedly and distinctly altered reward integration in both circuits. In mPFC-NAc (Fig. 4G), encoding of prior outcome was erased and only the current outcome encoded (Fig. 4H). This differs from both the two-lever and one-lever tasks wherein mPFC-NAc encoded a graded representation of reward history and suggests that mPFC-NAc integrates reward history only in instrumental settings where a response elicits outcomes. However, even when rewards are passively encountered (i.e. when no lever press is required), mPFC-NAc continues to encode reward but with a shortened time constant, such that only the most recent outcome is retained. vHip-NAc representation of reward history was also degraded, yet in a distinct manner (Fig. 4I). Current outcomes were encoded only when the previous trial, t-1, was unrewarded, suggesting a dependence on loss (Fig. 4J). This shift in encoding translates into vHip-NAc effectively overlooking isolated instances of non-reward, likely reflecting an extended time constant.

Critically, this cannot be explained by changes in task engagement given that mPFC-NAc continued to represent reward in these same animals (Fig. 4G,H) and licking bouts were similarly maintained across task variants (Fig. S5). Rather, this reveals that task demands differently shape neural encoding of reward in mPFC-NAc and vHip-NAc. When reward is passively encountered, independent of a required response, mPFC-NAc maintains a simplified reward representation across a shortened temporal window, limiting integration across trials. In contrast, vHip-NAc anchors encoding to loss with an extended time constant, to preferentially represent surprising rewards. This suggests that while the fundamental function of mPFC-NAc in rewarding contexts is to encode outcomes, the fundamental function of vHip-NAc is to use loss to tune outcome encoding.

### mPFC-NAc and vHip-NAc modulate task engagement

Examining neural representation of outcomes identified both mechanistic redundancy and functional specificity in mPFC-NAc and vHip-NAc encoding. We then asked how this neural processing might integrate to modulate behavior. While in general encoding was similar in both circuits, reducing the requirement for engagement by making reward non-contingent revealed functional specialization. We hypothesized that outcome-associated neural activity in mPFC-NAc and vHip-NAc modulates task engagement. To test this, we examined if neural activity at ITI end predicted latency to lever press on the subsequent trial, a metric operationalizing engagement (Bari et al., 2019; Beierholm et al., 2013; Cox et al., 2023; Hamid et al., 2016; Niv et al., 2007). A linear mixed effects model revealed modest yet significant relationships between latency to lever press and mPFC-NAc, vHip-NAc, and the interaction of mPFC-NAc and vHip-NAc activity (Fig. 5A, Supplementary Table 2; Fig. S6). This suggests that increased activity during outcome integration in either circuit increases latency to lever press, indicating reduced behavioral engagement.

**Figure 5.**
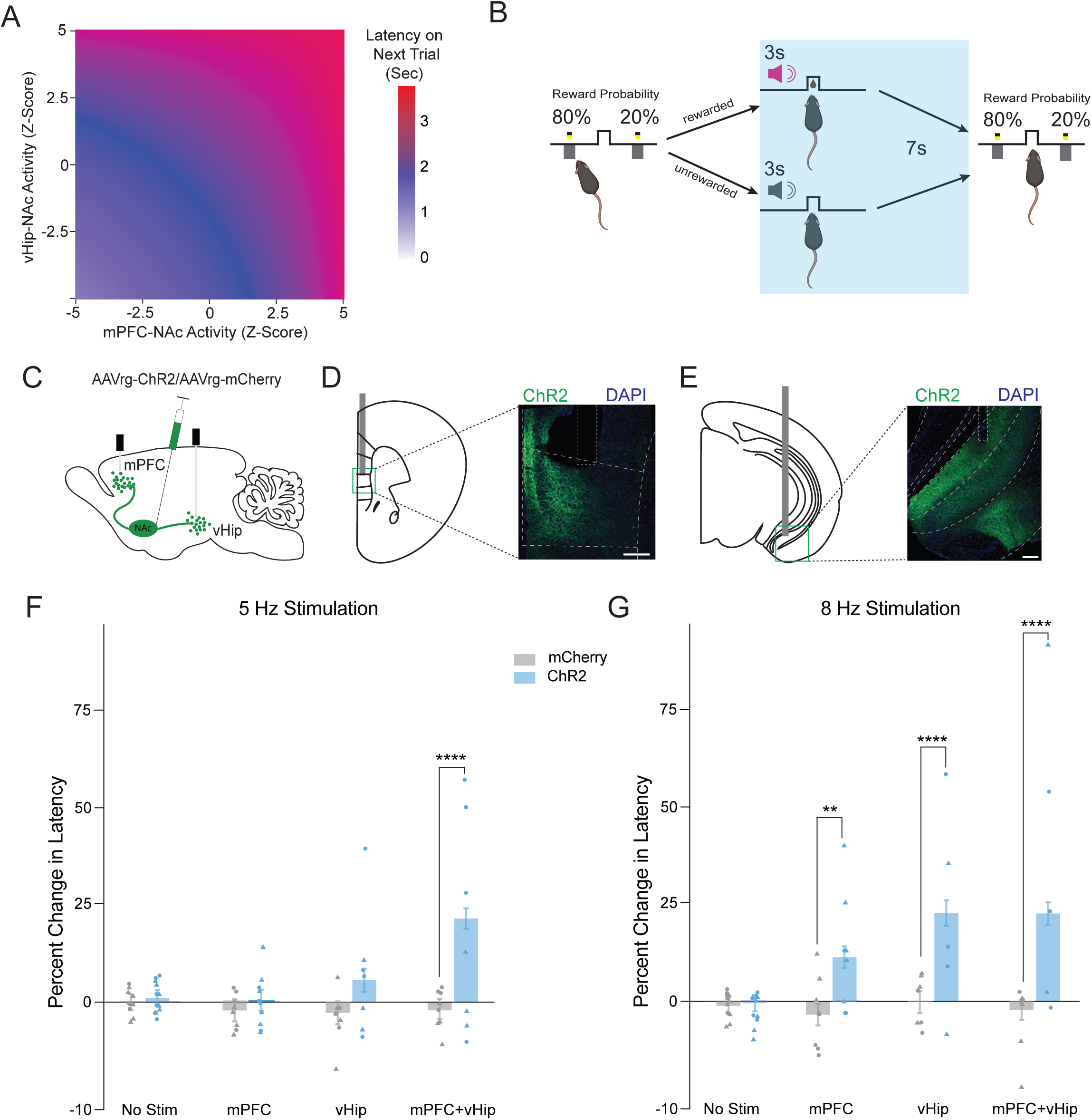
mPFC-NAc and vHip-NAc modulate task engagement. (A) Heatmap of estimated latency to respond on the subsequent trial given mPFC-NAc and vHip-NAc activity at ITI end shows that increased activity associates with longer latency. (B) Optogenetic stimulation in the two-armed bandit task is delivered for the duration of the ITI to either mPFC-NAc, vHip-NAc, or simultaneously to both circuits. (C) AAVrg-ChR2-mCherry or AAVrg-mCherry is injected into the NAc and optic fibers implanted in mPFC and vHip to stimulate (D) mPFC-NAc neurons and (E) vHip-NAc neurons. (F) Simultaneous 5Hz stimulation of mPFC-NAc and vHip-NAc, but neither circuit individually, increased latency to respond in ChR2 animals (male n=6, female n=7) compared to mCherry controls (male n=6, female n=6; Z=-18.6984, p<0.0001). (G) 8 Hz stimulation of mPFC-NAc (Z=-12.6970, p=0.01354), vHip-NAc (Z=-23.8073, p<0.0001), and simultaneous stimulation of both mPFC-NAc and vHip-NAc (Z=-24.1357, p<0.0001) all increased latency in ChR2 animals (male n= 5, female n=6) compared to mCherry controls (male n= 6, female n=6). Individual-animal averages are indicated by circles for males and triangles for females. Error bars represent SEM around the estimated mean. **p<0.01,****p<0.0001

From the association between neural activity and latency, we hypothesized that reward suppresses activity in mPFC-NAc and vHip-NAc to support behavioral engagement, defining a mechanism whereby recent reward history modulates engagement in reward-motivated behavior. We predicted that acutely increasing activity in either mPFC-NAc or vHip-NAc would suppress engagement. To test this, we injected retrograding AAV-ChR2 into NAc and implanted fibers above mPFC and vHip to deliver blue light stimulation during the ITI on a subset of trials in the two-armed bandit task (Fig. 5B-E; Fig. S7). To test if mPFC-NAc and vHip-NAc uniquely or redundantly control behavior we stimulated each circuit alone or both simultaneously.

Stimulating either circuit alone had no effect, whereas stimulating both simultaneously increased latency to lever press, but did not alter choice behavior (Fig. 5F; Fig. S8A). This could indicate either a threshold for sufficient cumulative glutamatergic drive or a requirement for synergistic interaction between inputs. To differentiate these possibilities, we repeated the experiment with stronger stimulation. Strong stimulation of either circuit alone increased latency to lever press, again with no effect on choice (Fig. 5G, Fig. S8B). This shows that total glutamatergic input modulates engagement, independent of input identity. mPFC-NAc stimulation yielded a slightly weaker effect than vHip-NAc, consistent with previous findings that mPFC projections to NAc medial shell are sparser than those from vHip (Britt et al., 2012). Stimulation during lever presentation did not yield any changes in latency or choice behavior, supporting the importance of neural integration of outcome during the ITI period, prior to action initiation (Fig. S9). Together our results demonstrate that mPFC-NAc and vHip-NAc dynamically track outcome information to modulate behavioral engagement according to recent history of reward. While each circuit is specialized to execute this function under distinct behavioral states, once engaged, they redundantly modulate behavior pointing to complementary roles in control of reward seeking.

## DISCUSSION

We examined redundancy and specificity in the function of two distinct glutamatergic inputs to the NAc. Using dual-site fiber photometry to probe trial-by-trial outcome encoding simultaneously in two circuits in the same animal during reward-guided choice, we find that mPFC-NAc and vHip-NAc similarly integrate reward via suppression of neural activity. By then systematically manipulating the conditions in which outcomes are encountered, we revealed that each circuit executes this common function under distinct behavioral states. While the mPFC-NAc invariantly encodes outcome, vHip-NAc uses information about loss to tune outcome encoding, effectively amplifying surprising reward. By comparing independent or synchronous circuit-specific optogenetic stimulation we show that, once engaged, these circuits cooperatively execute a shared function, i.e. modulating task engagement. Taken together, we identify a redundant mechanism for outcome integration with circuit-specific gating. This supports convergence of multiple inputs in tuning behavioral engagement to recent history of reward.

Our finding that both mPFC-NAc and vHip-NAc integrate information about outcomes of reward-motivated actions is consistent with the well-established role of mPFC in reward processing. Critically, we demonstrate that this function is not specific or limited to the mPFC-NAc.

Globally, the mPFC encodes information about previous actions and outcomes (Sul et al., 2010) and mPFC projections to the NAc bridge information about current actions and outcomes across trials (Parker et al., 2022; Spellman et al., 2021). Our findings suggest these functions are not unique to mPFC-NAc and are shared by vHip-NAc. However, we identify novel state dependent specialization in how reward integration is engaged in each circuit. We show that the mPFC-NAc fundamentally functions as a reward ledger, with reward suppressing neural activity no matter the behavioral state. In contrast, we find that vHip-NAc is tuned to preferentially encode outcome information after loss.

Differential encoding between mPFC-NAc and vHip-NAc emerged upon degrading task requirements, a manipulation that minimizes cognitive and behavioral demands effectively reducing the behavioral utility of representing integrated reward history. Under these circumstances, the base functionality of each circuit is revealed: mPFC-NAc encoding is anchored to reward whereas vHip-NAc is anchored to loss. Layered on top of this base functionality, representation of reward history scales with task complexity in support of behavioral demands. When reward is passively encountered with limited utility for action-outcome associations, mPFC-NAc encoding is limited to the most recent outcome. In more complex environments wherein actions elicit reward and action-outcome associations have high utility, the mPFC-NAc encoding window extends to integrate reward history. In simpler task structures that no longer require active engagement with a lever to earn rewards, the time-constant of vHip-NAc encoding shifts such that an activity no longer increases when a single unrewarded outcome follows reward. As a result, the vHip-NAc effectively comes to encode consecutive loss against other outcomes. Together, this suggests a role for vHip-NAc in providing information about the state of reward statistics in the environment, modulating behavior as a function of loss, and revealing a novel role for this circuit as a parallel and distinct stream of outcome integration.

The NAc has long been implicated in reward processing yet the precise neural circuit mechanisms are still being resolved. In the NAc medial shell, reward predominantly suppresses neural activity (Chen et al., 2023). This suppression likely maintains reward seeking as stimulation of either D1 or D2 medium spiny neurons bidirectionally controls reward seeking behavior (Lafferty et al., 2020). Here we show that reward suppresses both mPFC-NAc and vHip-NAc, two major excitatory inputs to NAc medial shell. Reward-associated suppression of these inputs would lead to reduced NAc activity. As such, our findings are consistent with reports that optogenetic stimulation of diverse glutamatergic inputs inhibits motivated behavior and the idea that glutamatergic input to NAc medial shell functions as a brake on motivated behavior (Lafferty et al., 2020; Millan et al., 2017; Reed et al., 2018; Yoshida et al., 2019, 2021). We show that outcome integration in mPFC-NAc and vHip-NAc initiates parallel, temporally integrated, neural signaling that may engage this ‘brake’ to align ongoing behavior with recent reward history and so tune behavioral engagement to prevailing environmental conditions.

Employing a redundant mechanism in mPFC-NAc and vHip-NAc may serve several functions. A common mechanism makes for simple integration of multiple inputs and ensures the robustness of the fundamental function of reward-guided engagement against insults. Further, modulating redundant encoding with state-dependent circuit-specific sensitivity may increase the granularity and range of encoding to ultimately amplify the behavioral impact of surprising rewards. We demonstrate that high levels of reward suppress activity in both mPFC-NAc and vHip-NAc to favor continued engagement. In contrast, strong activation of either input suppresses engagement, but, when weakly activated, synchronous recruitment of both circuits is required. Functionally, this may translate into a mechanism whereby moderate, balanced activity predominantly modulates task engagement while allowing for strong activation of either circuit to exert more direct behavioral control.

Preferential outcome encoding in vHip-NAc after loss may serve to strengthen engagement in variably rewarding environments, driving increased engagement when reward is infrequently encountered. The sensitivity of vHip-NAc to continuous loss, may also serve to gauge reward statistics of the environment, continually increasing with each consecutive loss to trigger task disengagement when activity reaches some threshold. Qualitatively, we see suggestion of this in the shape of the signal after loss: mPFC-NAc tends to plateau while vHip-NAc continues to increase. Ultimately, dysregulated outcome-encoding in either mPFC-NAc or vHip-NAc could alter behavioral sensitivity to reward and loss. Relative to mPFC-NAc, The vHip-NAc is poised to exert an outsized effect on behavioral engagement both in the strength of its input to NAc medial shell (Britt et al., 2012) and in its role in signaling loss. For example, hyperactivity of vHip-NAc may erroneously signal sustained loss, causing premature disengagement. Given our finding that engagement is modulated by the cumulative glutamatergic input to NAc, a sufficiently strong vHip-NAc signal could effectively jam any reward signal from mPFC-NAc, compounding insensitivity to reward that manifests as anhedonia. Indeed, disruption of the balance between NAc inputs and increased vHip-NAc drive is observed following chronic stress (Bagot et al., 2015; Muir et al., 2020; Pignatelli et al., 2021; Williams et al., 2020), as well as chronic alcohol (Griffin et al., 2023; Kircher et al., 2019) and cocaine intake (Barrientos et al., 2018; Cahill et al., 2016; Pascoli et al., 2014; Zinsmaier et al., 2022), manipulations associated with aberrant reward processing.

Here we examined the simultaneous encoding in two key neural circuits for motivated behavior. By considering outcome encoding within the context of recent outcome history and behavioral demands we identified a common neural mechanism of sustained temporal integration of reward outcomes and reveal how the external environment differentially shapes internal representations within two neural circuits. We also revealed critical circuit specificity: while mPFC-NAc consistently tracks outcomes, vHip-NAc preferentially encodes outcome information after loss. By illustrating the interplay of redundancy and specificity in circuit control of motivated behavior we demonstrate the need to contextualize events within varied behavioral states to fully understand neural encoding. Overall, our findings point to the importance of balanced suppression of NAc glutamatergic inputs during outcome integration to maintain reward-modulated behavioral engagement.

## METHODS

### Animals

Mice were maintained on a 12-h light-dark cycle (lights on at 7:00AM) at 22-25⁰C, group-housed with 3-4 same-sex cage-mates with *ad libitum* access to food and water. All experimental manipulations occurred during the light cycle, in accordance with guidelines of McGill University’s Comparative Medicine and Animal Resources Center and approved by the McGill Animal Care Committee. 7-week-old male and female C57BL/6J mice were obtained from Jackson Laboratories and habituated to the colony room one week prior to start of manipulations. Mice were food restricted to 85% of their free-feeding body weight during experimentation.

### Surgeries

Stereotaxic surgery was performed under ketamine (100 mg/kg)/xylazine (10 mg/kg) anesthesia. To achieve projection-specific GCaMP7f expression in glutamatergic NAc-projecting cells, 0.3μl pGP-AAVrg-syn-jGCaMP7f-WPRE virus (1.85× 10^13^GC/ml; Addgene) was infused into the NAc (A/P: +1.3, M/L: +/-0.60, D/V: −4.9) at a rate of 0.1μl per min, before raising the needle to D/V: −4.7 and infusing a further 0.4µl virus, and allowed to diffuse for 10 min before withdrawing the needle. pGP-AAV-syn-jGCaMP7f-WPRE was a gift from Douglas Kim & GENIE Project (Addgene plasmid # 104488; http://n2t.net/addgene:104488; RRID:Addgene_104488) (Dana et al., 2019). Chronically implantable optic fibers (Neurophotometrics) with 200μm core and 0.37 NA threaded through ceramic ferrules were implanted above the ventral subiculum of the vHip (A/P: −3.40, M/L: +/-3.00, D/V: −4.75) and infralimbic mPFC (A/P: 1.90, M/L: +/-0.3, D/V: −2.80). Recordings began minimum 4 weeks after surgery to allow sufficient time for stable and robust retrograde virus expression. To achieve projection-specific ChR2 expression in glutamatergic NAc-projecting cells, 0.3μl pGP-AAVrg-hSyn-hChR2(H134R)-EYFP virus (7× 10^12^GC/ml; Addgene) or a fluorophore only control, pGP-AAVrg-hSyn-mCherry (7× 10^12^GC/ml; Addgene) was infused into the NAc (A/P: +1.3, M/L: +/-0.60, D/V: −4.9) at a rate of 0.1μl per min, before raising the needle to D/V: −4.7 and infusing a further 0.4µl virus, and allowed to diffuse for 10 min before withdrawing the needle. pAAV-hSyn-hChR2(H134R)-EYFP was a gift from Karl Deisseroth (Addgene plasmid # 26973; http://n2t.net/addgene:26973; RRID:Addgene_26973). pAAV-hSyn-mCherry was a gift from Karl Deisseroth (Addgene plasmid # 114472; http://n2t.net/addgene:114472; RRID:Addgene_114472). Chronically implantable optic fibers (Neurophotometrics) with 200μm core and 0.22 NA threaded through ceramic ferrules were implanted above the ventral subiculum of the vHip (A/P: −3.40, M/L: +/-3.00, D/V: −4.75) and infralimbic mPFC (A/P: 1.90, M/L: +/-0.3, D/V: −2.80). Optogenetic manipulations began minimum 4 weeks after surgery to allow sufficient time for stable and robust retrograde virus expression.

### Histology

After completion of all behavioral testing, mice were deeply anesthetized with ketamine/xylazine and transcardially perfused with phosphate buffered saline (PBS) and paraformaldehyde (4%). Brains were removed and post-fixed in paraformaldeyhde for 24h and stored in PBS until sectioning on a vibratome (50 µm). Sections were mounted with Vectashield with DAPI (Vector Laboratories) and examined under a fluorescent microscope (Leica DM6000 B) to confirm viral expression and fiber placement. A confocal microscope (Zeiss LSM800) was used to obtain fluorescent images. Images were acquired as tiles with a 20x air objective (NA 0.8) using Zeiss Zen Blue imaging software. Images were collected in the McGill University Advanced BioImaging Facility (ABIF), *RRID:SCR_017697*. Mistargeted animals were excluded from analysis.

### Apparatus

Behavioral experiments were performed in standard Med Associates operant boxes (15.24 x 13.34 x 12.7 cm) enclosed in sound attenuating chambers outfitted with a programmable audio generator, two retractable levers and cue lights either side of a food port for delivering a liquid chocolate milk reward (30μl, Nesquick) diluted with water in a 2:1 ratio. Boxes were controlled and data collected by a computer running MED-PC software (Med-Associates).

### Lever Press Training

Training was completed in three stages, with all training sessions lasting 30 minutes. In the first stage, animals were presented with two levers, both of which delivered a chocolate milk reward with a 100% probability. To signal the start of the trial, both levers extended and the cue lights above the levers turned on, animals then had 60 seconds to make a response on either lever. A press on either lever resulted in lever retraction, immediate delivery of a 30 µL chocolate milk reward, and the start of a 3 second auditory cue (2kHz pure tone or white noise). Following either a lever press or 60 seconds with no press (i.e. an omission), a 10 second intertrial interval (ITI) was triggered. After one session with over 25 responses, animals progressed to the second stage. In this stage animals again were presented with two levers but reward was now delivered with a 50% probability on both levers. To signal the start of the trial, both levers extended and the cue lights above the levers turned on, animals then had 60 seconds to make a response. A lever press resulted in lever retraction and immediate delivery of the outcome, either a 30 µL chocolate milk reward and a 3 second auditory cue (2kHz pure tone or white noise, counterbalanced across animals) or just a 3 second auditory cue (white noise or 2kHz pure tone). Following either a lever press or omission, a 10 second intertrial interval (ITI) was triggered. Following two consecutive sessions with over 40 responses, animals progressed to the third stage. This stage was the same as stage two except that animals now had only 10 seconds to make a response before an omission was registered. Following two consecutive sessions with over 100 responses animals achieved criterion to progress to the two-armed bandit task.

### Two-armed bandit Task

The two-armed bandit task was performed over of 6 days with each session lasting one hour. In this task, animals were presented with two levers with one lever rewarded on 80% of trials, and the other lever rewarded on 20% of trials. To signal the start of the trial, both levers extended and the cue lights above the levers turned on, animals then had 10 seconds to make a response on either lever or an omission was registered. A lever press resulted in lever retraction and immediate delivery of the outcome, either a 30 µL chocolate milk reward and a 3 second auditory cue (2kHz pure tone or white noise, counterbalanced across animals) or simply a different 3 second auditory cue (white noise or 2kHz pure tone) signaling non-reward. Following either a lever press or an omission, a 10 second intertrial interval (ITI) was triggered. To maintain a dynamic learning environment and high rates of rewarded and unrewarded outcomes, probability of reward was switched between levers after five consecutive responses on the high probability lever. Four males and four females remained on the two-armed bandit task during the task degradation (data not shown).

### One-Lever Forced Choice Task

The one lever forced choice task was performed over 3 days with each session lasting one hour. In this task, animals were presented with a single lever (counterbalanced across animals). Pressing this lever resulted in probabilistic reward on a predetermined schedule. The outcome schedule was matched to each animal’s individual performance in the final three days of the two-armed bandit task, such that the first session in the one-lever task was yoked to the reward schedule experienced by the animal on day four in the two-armed bandit task, the second to day five, and the third to day six. To signal the start of the trial, the lever extended and the cue light above the lever turned on. Animals then had 10 seconds to make a response. A lever press resulted in lever retraction and immediate delivery of the outcome, either a 30 µL chocolate milk reward and a 3 second auditory cue (2kHz pure tone or white noise) or simply a different 3 second auditory cue (white noise or 2kHz pure tone). Following either a lever press or an omission, a 10 second intertrial interval (ITI) was triggered.

### No Lever Response Free Task

The no lever response free task was performed over the course of 3 days with each session lasting one hour. In this task, animals were able to retrieve non-contingently delivered rewards under a similar trial structure to both the two-armed bandit task and the one-lever forced choice task but with no levers available. To signal the start of the trial, cue lights above both levers turned on and remained illuminated for a period of time matched to each animal’s response time in the last three days of the two-armed bandit task. After cue lights turned off, outcomes were delivered, either a 30 µL chocolate milk reward and a 3 second auditory cue (2kHz pure tone or white noise) or simply a different 3 second auditory cue (white noise or 2kHz pure tone). As in the one-lever task, the outcome schedule was matched to each animal’s performance in the final three days of the two-armed bandit task now also matching the latency to receive the outcome to the trial-by-trial latency to lever press on the two-armed bandit task with a 10 second intertrial interval (ITI).

### Frame Independent Projected Fiber Photometry

To measure calcium-associated changes in fluorescence in real time, recordings were made from vHip-NAc and mPFC-NAc-projecting cells during the two-armed bandit task, the one-lever forced choice task, and the no lever response free task. Samples were collected at a frequency of 20 Hz using Neurophotometrics hardware through Bonsai and FlyCap software. Recordings were coupled to the start of behavioral analysis by interfacing Bonsai with MED-PC using a custom DAQ box (Neurophotometrics).

### Photometry data extraction and normalization

*Photometry data* were extracted and analyzed using custom-written scripts in Python. To normalize the data, the control channel (415nm) was fitted to the raw (470nm). The fitted control was then subtracted from the raw trace. The resultant trace was divided by the fitted control giving the ΔF/F and converted to a Z-score. This calculation was performed over the entirety of the session to preserve dynamic fluctuation in population activity that persists beyond individual trials to allow comparison across trials. For heatmaps Z-scores were baseline subtracted from average activity in the two seconds prior to lever press to accommodate moving baselines. For analyses of reward history, Z-scores were baseline subtracted from average activity in the two seconds prior to lever press on trial t-1 to account for shifted baselines in trial t0.

### Optogenetics in Two-armed Bandit Task

Following lever press training, animals started the two-armed bandit task with optogenetic manipulations of mPFC-NAc and vHip-NAc activity for the duration of the ITI. Each day animals received either mPFC-NAc, vHip-NAc, or simultaneous mPFC-NAc and vHip-NAc stimulation on a subset of trials over the course of 9 days such that they received a total of 3 days of stimulation per condition for each stimulation protocol tested (5 Hz, 10 ms, 1-2 mW; 8 Hz, 10 ms, 2-3 mW). Order of stimulation days was fully counterbalanced within and between mice to avoid any order effects. Stimulation was delivered by 450 nm lasers controlled by a laser driver (Doric) running Doric studios software and triggered via a TTL (Med-Associates) at ITI start on a random subset of trials (30%) and terminated immediately prior to lever extension.

### Ex vivo current-clamp electrophysiology

Brain slice preparation

Mice were deeply anesthetized with isofluorane. Transcardial perfusion was performed with 25-30 ml of ice-chilled carbogenated NNMDG artificial cerebrospinal fluid (aCSF: containing in mM: 92 NMDG, 2.5 KCl, 1.25 NaH_2_PO_4_, 30 NaHCO_3_, 20 HEPES, 25 glucose, 2 thiourea, 5 Na-ascorbate, 3 Na-pyruvate, 0.5 CaCl_2_·4H_2_O and 10 MgSO_4_·7H_2_O; titrated to pH 7.3–7.4 with concentrated hydrochloric acid). Brain slices (200 μm) were prepared in ice-chilled carbogenated NMDG aCSF by a vibratome (Lecia VT 1200S). All brain slices were recovery in 32–34 °C carbogenated NMDG aCSF for 10 min and then were transferred into room-temperature carbogenated HEPES holding aCSF (containing in mM: 92 NaCl, 2.5 KCl, 1.25 NaH_2_PO_4_, 30 NaHCO_3_, 20 HEPES, 25 glucose, 2 thiourea, 5 Na-ascorbate, 3 Na-pyruvate, 2 CaCl_2_·4H_2_O and 2 MgSO_4_·7H_2_O; titrated to pH to 7.3–7.4 with NaOH) for at least 1 hour before current-clamp recording.

Electrophysiology recordings

Current-clamp recordings were performed in room-temperature carbogenated aCSF (containing in mM: mM: 128 NaCl, 3 KCl, 1.25 NaH_2_PO_4_, 2 MgCl_2_, 2 CaCl_2_, 24 NaHCO_3_ and 10 glucose; pH 7.2). The patch pipette solution was composed of (in mM) 115 K-gluconate, 20 KCl, 1.5 MgCl_2_, 10 Phosphocreatine-Tris, 2 Mg-ATP, 0.54 Na-GTP and 10 HEPES. Blue light (wavelength: 470 nm) from a LED system (DC4100, Thorlabs) was used for optogenetic stimulation to evoke action potentials. The optogenetic stimulation protocol consisted of trains of 5 Hz (1-2 mW) or 8 Hz (2-3 mW) 10ms light pulses for 5 s. All signals were amplified and digitized by Multiclamp 700B (Molecular Device) and Digidata 1550B (Molecular Device) respectively. Series and access resistance were monitored during the experiments and signals were bessel filtered at 2 kHz.

### Data Analysis & Statistics

#### Linear Mixed Effects Regression

Linear Mixed Effects Regression Models are a powerful approach to probe variance attributable to variables of interest (e.g. trial outcome) while simultaneously controlling for random effects (e.g. session ID) (Fetcho et al., 2023; Kato et al., 2022; Yu et al., 2022). This is useful for modeling instances where there is nonindependence in the structure of data e.g. multiple trials recorded within multiple animals. Models were fit using the full interaction of the factors of interest (trial outcome, previous trial outcome, sex) and using animal ID and session ID as random effects using the *lme4* package in R (Bates et al., 2014). Where the dependent variable was latency, a Gamma link function was used to approximate the non-gaussian distribution. The fitted models were used to calculate estimated marginal means using the *emmeans* package in R (Lenth et al., 2021). The effect of variables of interest were then examined by comparing estimated marginal means. Given the large number of samples generated using this approach (all trials x all animals), comparisons of estimated marginal means were conducted using a Z-test and Sidak’s method to adjust for multiple comparisons.

#### Cross-Correlation Time Delay Analysis

Time delay analysis was performed by first calculating the cross-correlation between mPFC-NAc and vHip-NAc during the ITI across a maximum lag of ± 5 seconds using the CCF function in R. The argument of the maximum (i.e. the time offset of peak correlation) of the resulting cross-correlation function was used to estimate the delay between mPFC-NAc and vHip-NAc on a trial-by-trial basis (Abboud & Sadeh, 1984). Linear mixed effects models were then fit to assess if the delay was non-zero (i.e. non-synchronous) using the following models to test for effects of sex [Time Delay∼Sex-1+(1|ID)+(1|Day)] and for the interaction between sex and reward [Time Delay∼Rewards:Sex-1+(1|ID)+(1|Day)]. The resulting regression coefficients from each model were examined to determine if the time delay was non-zero in any group (i.e. regression coefficient significantly different from zero).

#### Conditional Entropy Analysis

Conditional entropy is an information measure used to estimate the amount of additional information needed to explain one signal given full knowledge of a second signal. This can be interpreted as the unique information contributed by a second signal beyond that contributed by a first with smaller conditional entropy suggesting less unique information carried by the second signal. Conditional entropy was calculated on the first two seconds and the last two seconds of the ITI using the PyInform package in Python to calculate the entropy (□) of the mPFC circuit given the vHip-NAc circuit, □(mPFC-NAc|vHip-NAc), and the entropy of the vHip-NAc circuit given the mPFC-NAc circuit, □(vHip-NAc|mPFC-NAc) (Cover & Thomas, 1991; Moore et al., 2018).

## Code Availability

Code used to perform analyses for all figures available at https://github.com/eshaaniyer/mPFCvHip-NAc_RewardIntegration

## ACKNOWLEDGMENTS

We would like to thank Dr. Becket Ebitz, Dr. Mihaela Iordanova and Heike Schuler for their helpful comments and feedback throughout this project. Renderings of mice were created with BioRender.com. Research was supported by grants to RCB from NSERC and the Ludmer Centre for Neuroinformatics and Mental Health.

## Author contributions

Conceptualization, E.S.I and R.C.B.; Methodology, E.S.I and R.C.B.; Investigation, E.S.I, P.V., S.W., J.M., Y.C.T., V.C.; Writing – Original Draft, E.S.I and R.C.B.; Writing – Review & Editing, E.S.I. and R.C.B.; Funding Acquisition, R.C.B.; Resources, R.C.B.; Supervision, R.C.B.

