## Supplemental Materials for "Reward integration in prefrontal-cortical and ventral-hippocampal nucleus accumbens inputs cooperatively modulates engagement"

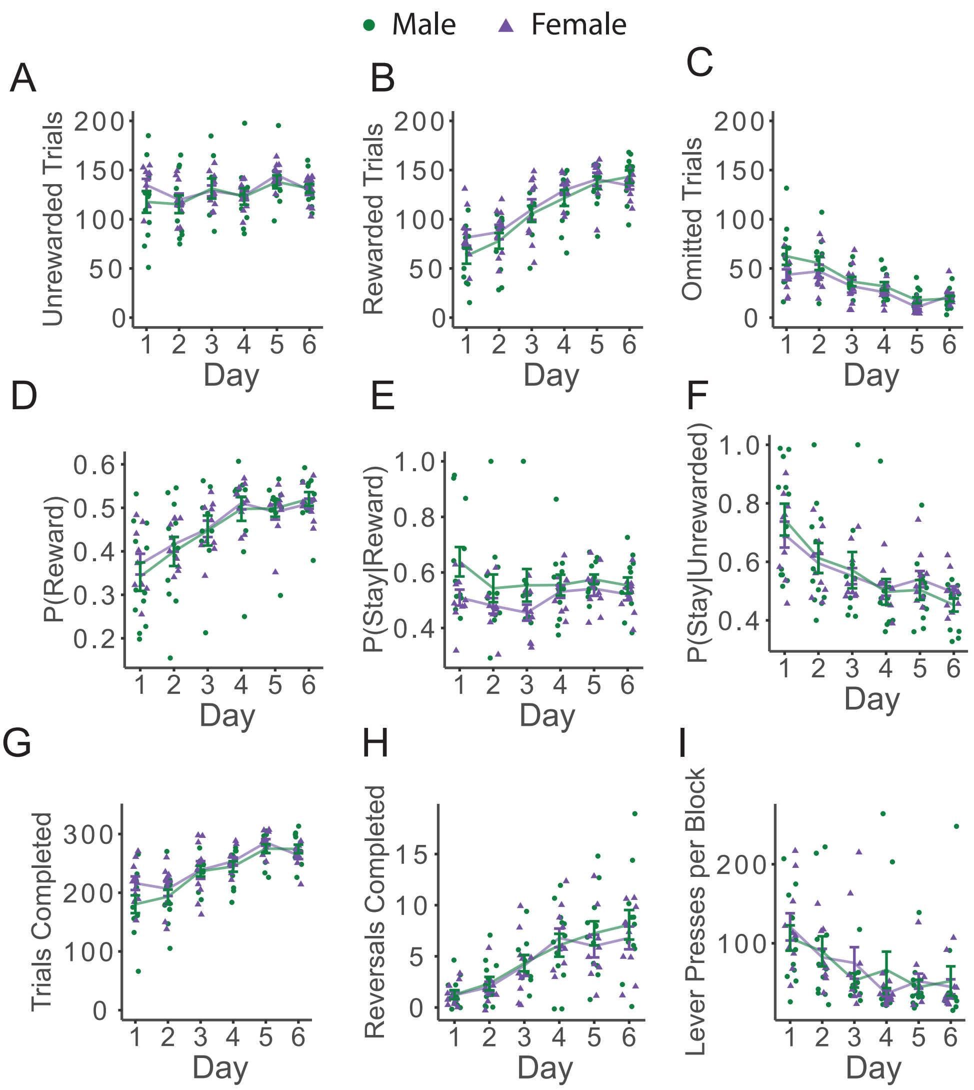


Supplementary Figure 1. Behavior in the two-arm bandit task across days. Male (n=12) and female (n=10) mice trend towards experiencing more (A) unrewarded trials across days (F=3.9919, p=0.0595, and experience more (B) rewarded trials across days (F=138.7239, p<0.0001), and fewer (C) omitted trials across days (F=51.5277, p<0.0001). (D) Across days, animals are more likely to earn rewards. Across days, animals do not change their staying probability following a (E) rewarded outcome but decrease their staying probability following (F) unrewarded outcomes (F=50.0712, p<0.0001). (G) Animals complete more trials across days (F=150.3925, p<0.0001), and increase the number of (H) reversals completed (F=40.3985, p<0.0001), and (I) decrease the number of lever presses performed to trigger a reversal (F=16.5760, p=0.0006). Individual-animals averages are indicated by circles for males and triangles for females. Error bars represent SEM around the estimated mean.

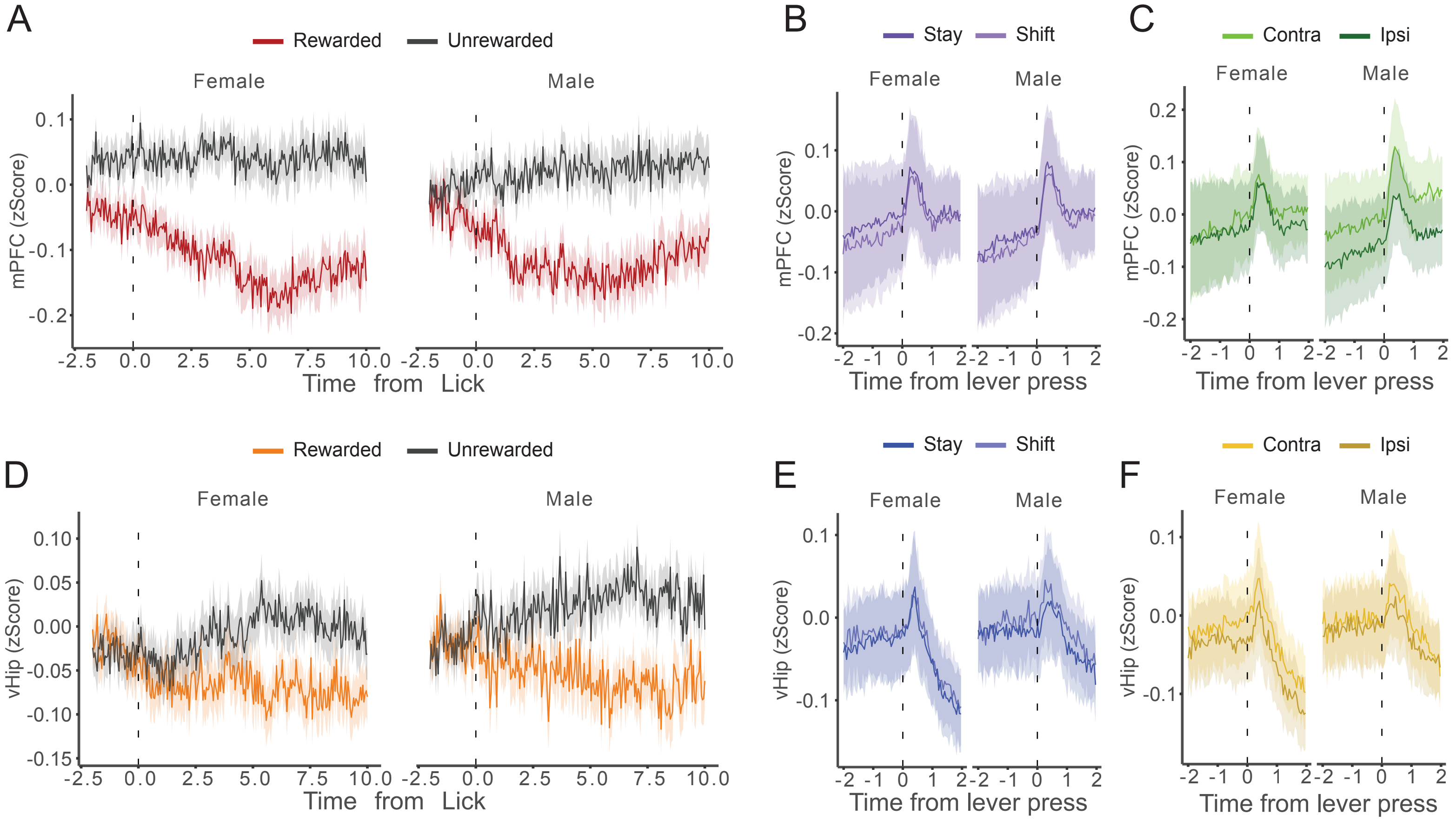


Supplemental Figure 2. mPFC-NAc and vHip-NAc activity time-locked to task-relevant features. (A) Estimated mean mPFC-NAc activity time-locked to the first rewarded and unrewarded lick during the ITI in female (n=10) and male (n=12) mice. (B) Estimated mean mPFC-NAc activity time-locked to lever press on trials in which the previous choice is repeated (stay) and when a different choice is made (shift) (C) Estimated mean mPFC-NAc activity time-locked to lever press on trails in which animals choose the lever contralateral or ipsilateral to their implant. (D) Estimated mean vHip-NAc activity time-locked to the first rewarded and unrewarded lick during the ITI in female (n=10) and male (n=12) mice. (E) Estimated mean vHip-NAc activity time-locked to lever press on trials in which the previous choice is repeated (stay) and when a different choice is made (shift) (F) Estimated mean mPFC-NAc activity time-locked to lever press on trails in which animals choose the lever contralateral or ipsilateral to their implant.

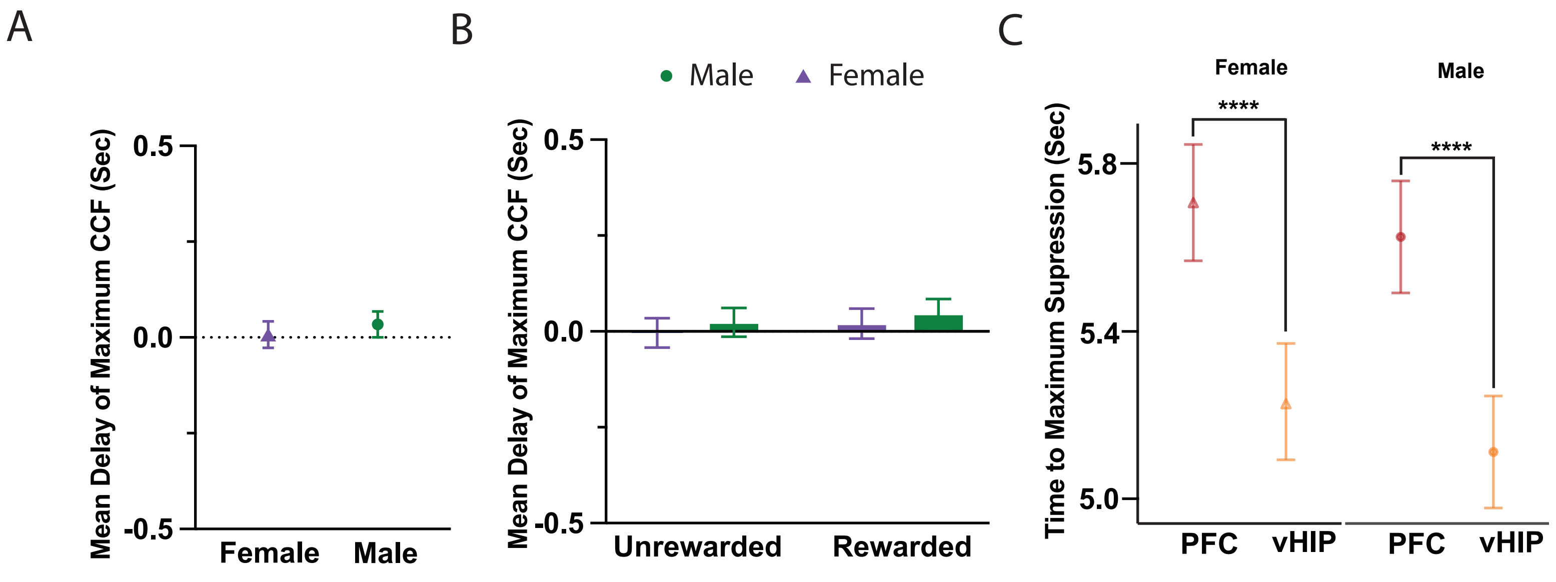


Supplementary Figure 3. Reward-associated encoding emerges similarly and is not led by either mPFC-NAc or vHip-NAc. (A) Regression coefficients for a linear mixed effects model predicting the time delay of the maximum cross-correlation do not significantly differ from zero in either females (n=10; Female=0.0074 ± 0.0349) or males (n=12; Male=0.03349 ± 0.0338). (B) Regression coefficients remain non-significant when analyzing cross-correlation separated by trial outcome (Female Unrewarded =-0.0035 ± 0.0382, Female Rewarded =0.0202 ± 0.0393, (Male Unrewarded =0.0237 ± 0.0373, Male Rewarded =0.0463 ± 0.0385). (C) Maximum suppression emerges at shorter latencies in vHip-NAc than mPFC-NAc in both females (Z=9.8839, $p<0.0001$) and males (Z=10.6193, $p<0.0001$). Error bars represent SEM around the estimated mean. ****$p<0.0001$

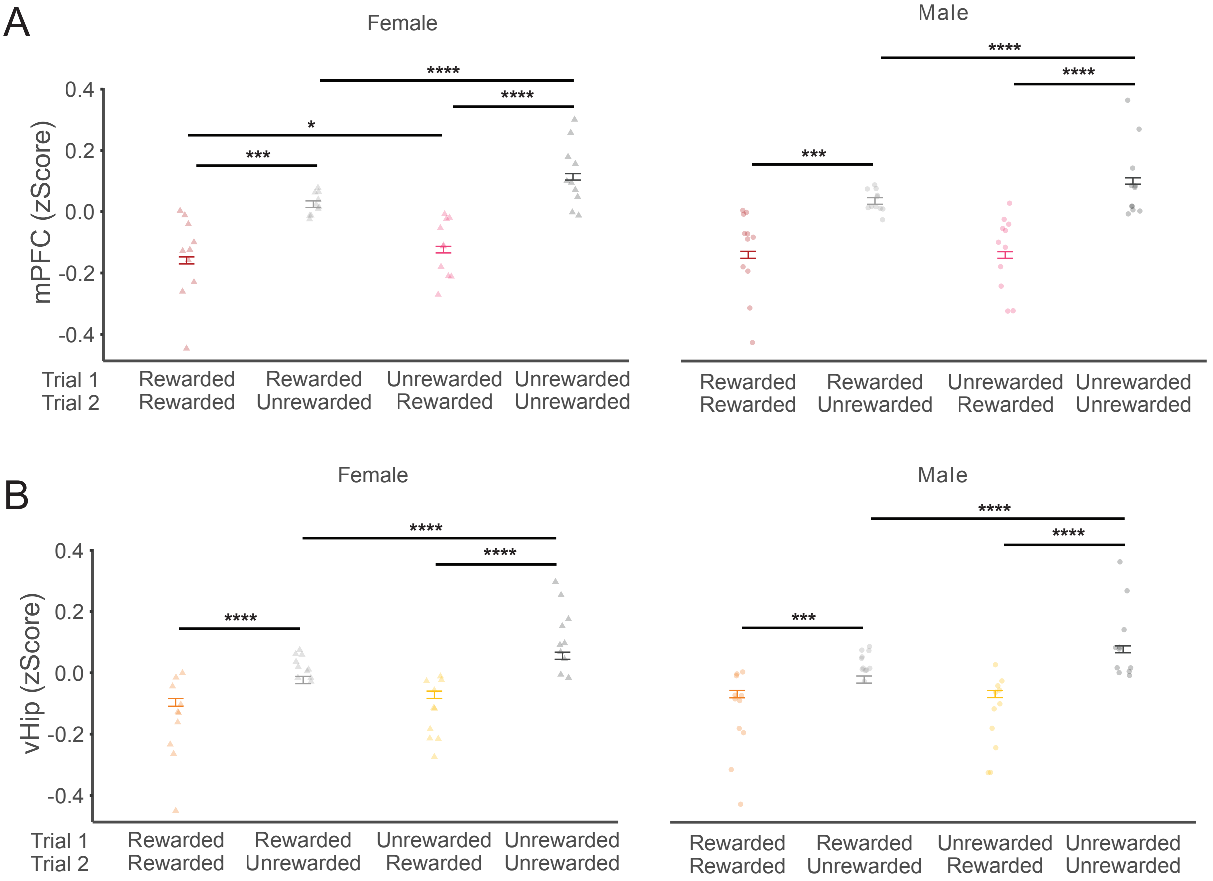


Supplementary Figure 4. mPFC-NAc and vHip-NAc similarly integrate reward history in females and males. Estimated mean neural activity across pairs of consecutive trials (t-1→t0) showing rewarded+rewarded (R→R), rewarded+unrewarded (R→U), unrewarded+rewarded (U→R), and unrewarded+unrewarded (U→U) trial pairs in female (n=10) and male (n=12) mice. Analysis focused on 8-10 sec after lever press (ITI end) on trial t0. (A) On trial t0, mPFC-NAc activity is significantly suppressed by current reward in females (U→U vs U→R: Z=-20.0296, $p<0.0001$; R→U vs R→R: Z=-14.4311, $p<0.0001$) and males ( U→U vs U→R: Z=-20.3516, $p<0.0001$; R→U vs R→R: Z=-13.7364, $p<0.0001$). When trial t0 is unrewarded, mPFC-NAc activity remains significantly suppressed by reward experienced on the previous trial, t-1, in females (U→U vs R→U: Z= 7.5245, $p<0.0001$) and males (U→U vs R→U: Z= 5.4925, $p<0.0001$). When trial t0 is rewarded, mPFC-NAc activity remains significantly suppressed by reward experienced on the previous trial, t-1, in females (U→R vs R→R: Z= 2.7695, $p=0.0223$). (B) On trial t0, vHip-NAc activity is significantly suppressed by current reward in females (U→U vs U→R: Z=-11.2415, $p<0.0001$; R→U vs R→R: Z=-6.0202, $p<0.0001$) and males U→U vs U→R: Z=-13.4823, $p<0.0001$; R→U vs R→R: Z=-4.0837, $p=0.0002$). When trial t0 is unrewarded, vHip-NAc activity remains significantly suppressed by reward experienced on the previous trial, t-1, in females (U→U vs R→U: Z= 6.9947, $p<0.0001$) and males (U→U vs R→U: Z= 9.1069, $p<0.0001$). Individual animal averages are indicated by circles for males and triangles for females. Error bars represent SEM around the estimated mean. *$p<0.05$, ***$p<0.001$, ****$p<0.0001$

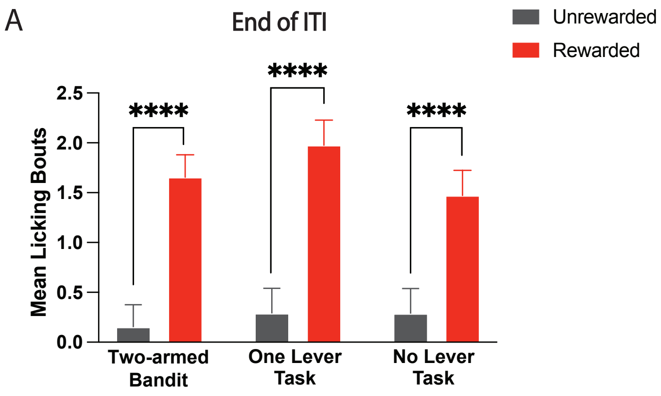


Supplementary Figure 5. Licking behavior is similar across task variants. (A) Licking bouts are similarly increased at ITI end on rewarded compared to unrewarded trials across task variants with no significant differences in licking behavior across task variants (male n= 8, female n=6; Two-lever Task: Z=-22.415, p<0.0001; One-lever Task: Z=-18.599, p<0.0001, No Lever Task: Z=-22.415, p=0.0001). ****p<0.0001

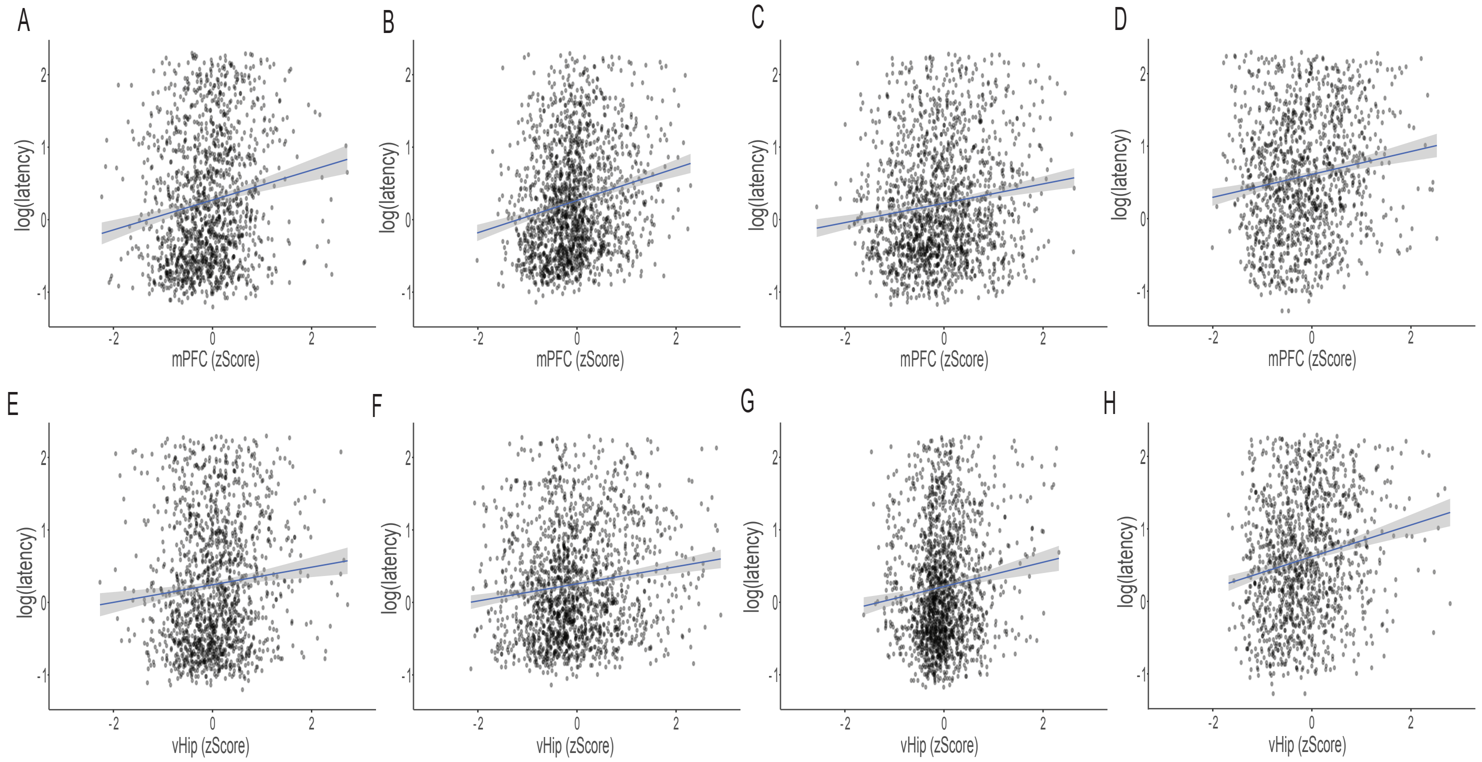


Supplemental Figure 6. Correlation between latency and mPFC-NAc or vHip-NAc activity at ITI end in representative animals. (A) Correlation between mPFC-NAc and latency at ITI end in animal #207 (male, $r=0.1542$, $p<0.0001$). (B) Correlation between mPFC-NAc and latency at ITI end in animal #208 (male, $r=0.1868$, $p<0.0001$). (C) Correlation between mPFC-NAc and latency at ITI end in animal #215 (female, $r=0.1368$, $p<0.0001$). (D) Correlation between mPFC-NAc and latency at ITI end in animal #215 (female, $r=0.1408$, $p<0.0001$). (E) Correlation between vHip-NAc and latency at ITI end in animal #207 (male, $r=0.1029$, $p<0.0001$). (F) Correlation between vHip -NAc and latency at ITI end in animal #208 (male, $r=0.1373$, $p<0.0001$). (G) Correlation between vHip-NAc and latency at ITI end in animal #215 (female, $r=0.1082$, $p<0.0001$). (H) Correlation between vHip -NAc and latency at ITI end in animal #215 (female, $r=0.1838$, $p<0.0001$).

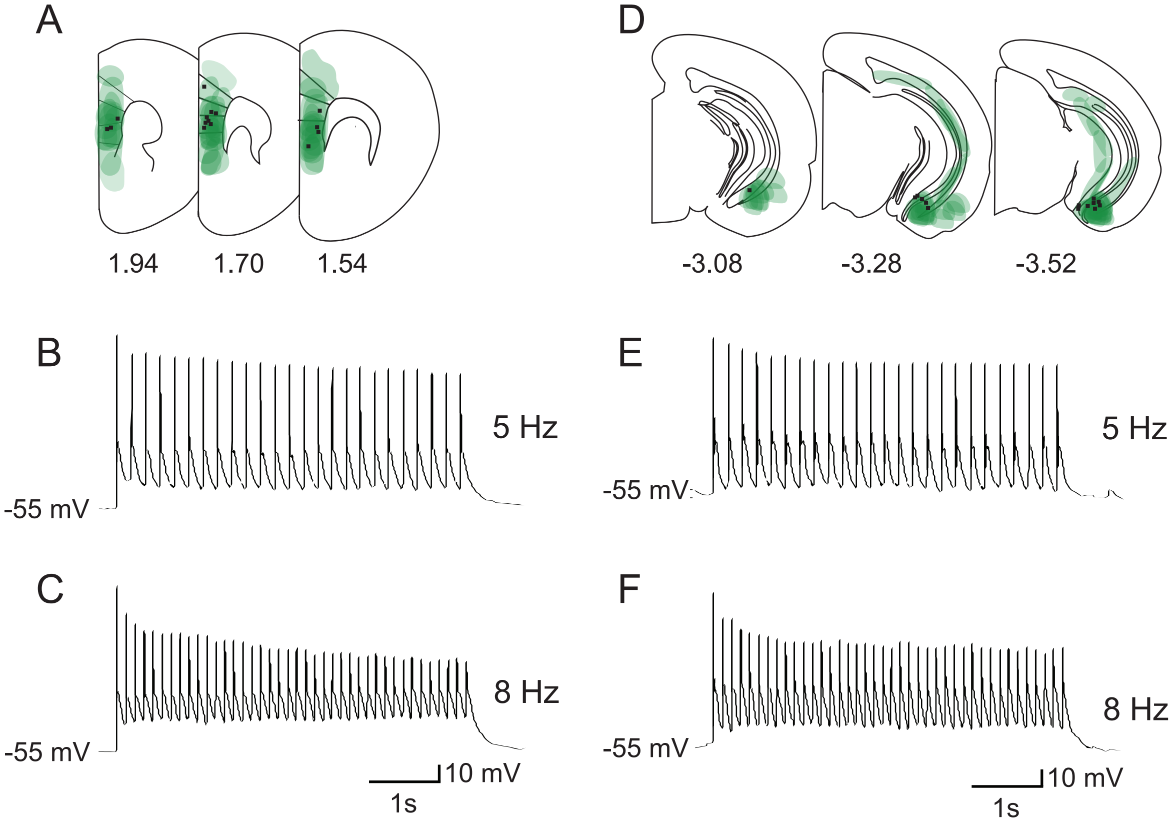


Supplementary Figure 7. Histology and optogenetic validation. (A) Plates indicate viral spread and location of fiber tips in the mPFC. In vitro optical stimulation of mPFC-NAc cell bodies induces spiking that reliably tracks stimulation at (B) 5Hz and (C) 8Hz. (D) Plates indicate viral spread and location of fiber tips in the vHip. In vitro optical stimulation of vHip-NAc cell bodies induces spiking that reliably tracks stimulation at (E) 5Hz and (F) 8Hz.

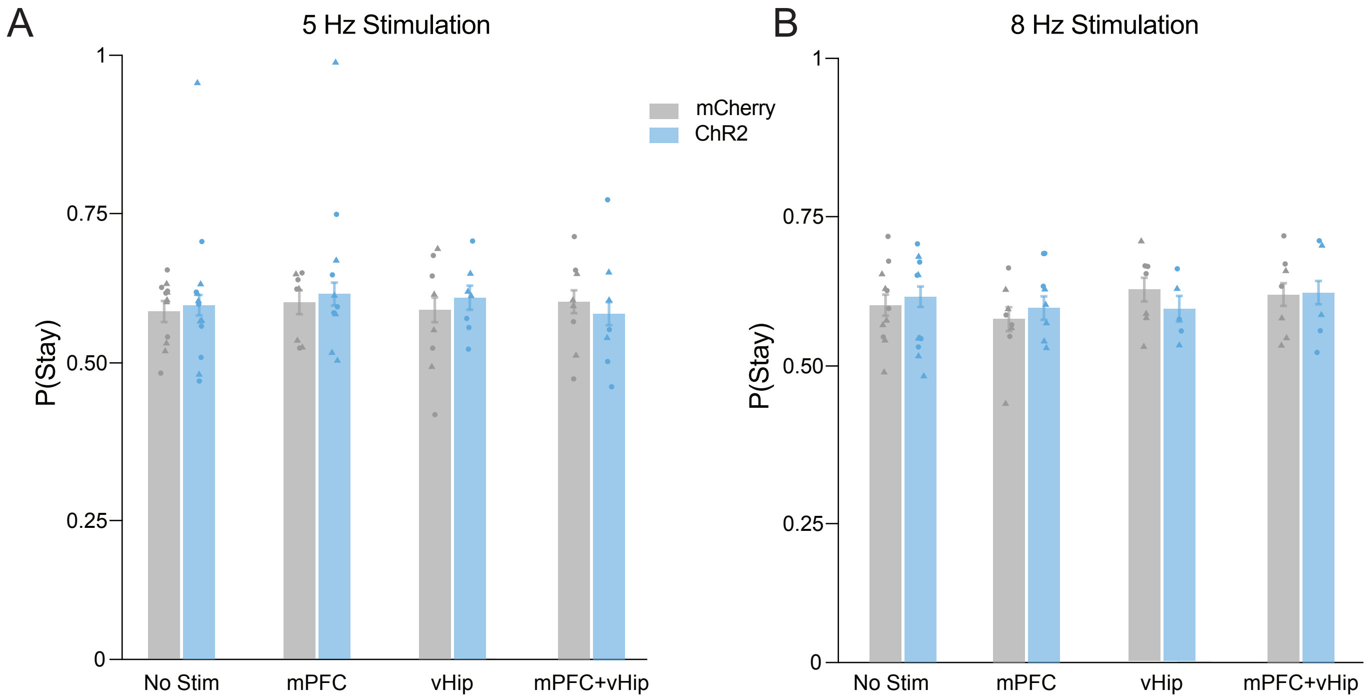


Supplementary figure 8. Optogenetic stimulation does not impact choice behavior. (A) Neither simultaneous nor individual 5Hz stimulation of mPFC-NAc and vHip-NAc changed staying probabilities in ChR2 animals (male n=6, female n=7) compared to mCherry controls (male n=6, n=6 females). (B) Neither simultaneous nor individual 8Hz stimulation of mPFC-NAc and vHip-NAc changed staying probabilities in ChR2 animals (male n=6, female n=7) compared to mCherry controls (male n=6, female n=6). Individual-animal averages are indicated by circles for males and triangles for females. Error bars represent SEM around the estimated mean.

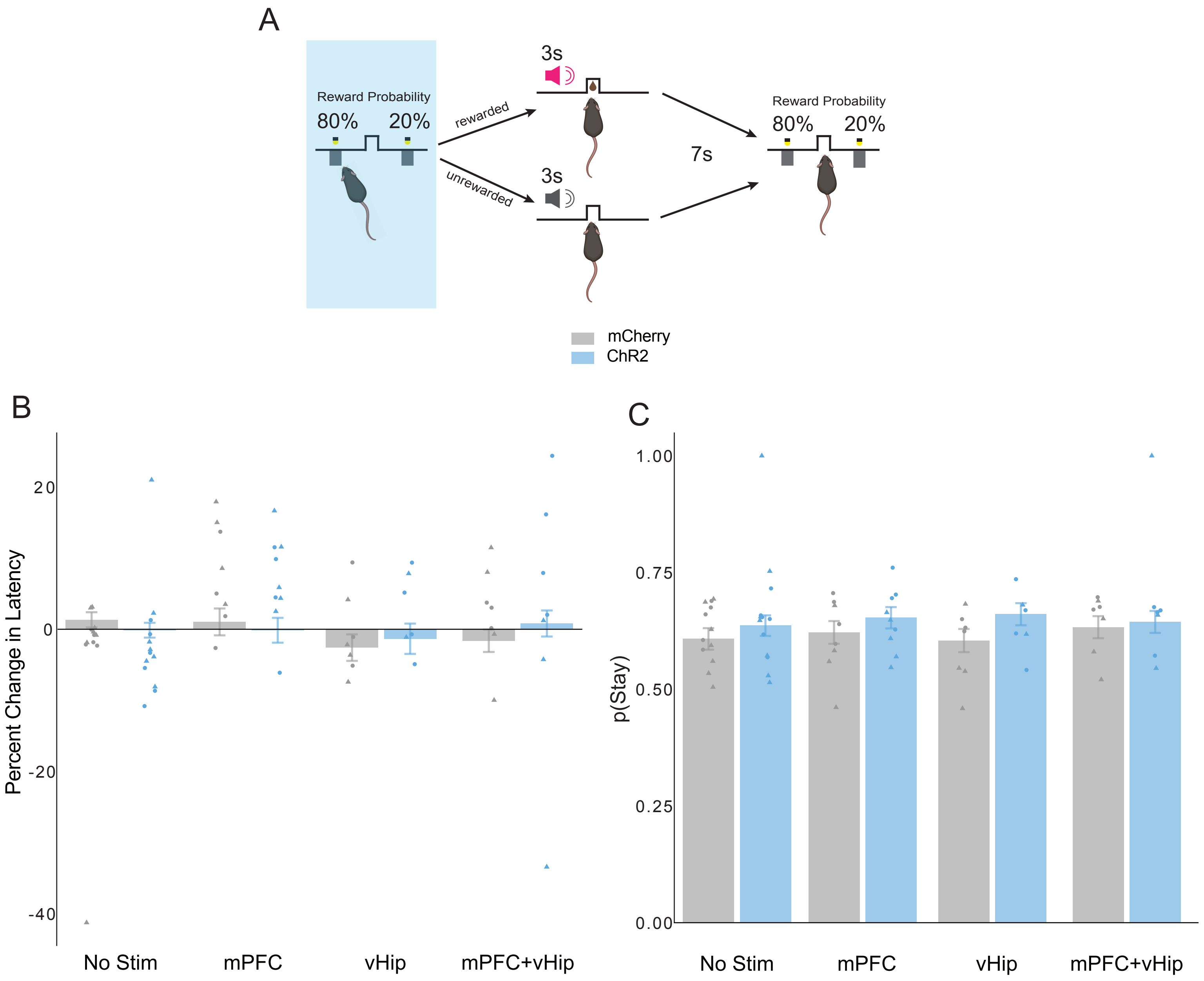


Supplementary figure 9. Optogenetic stimulation during lever-pressing does not impact behavior. (A) Optogenetic stimulation protocol in the two-armed bandit task is delivered for the duration of the lever extension to either mPFC-NAc, vHip-NAc, or simultaneously to both circuits. Neither simultaneous nor individual 5Hz stimulation of mPFC-NAc and vHip-NAc affected (B) latency to press or (C) staying probabilities in ChR2 animals (male n=6, female n=7) compared to mCherry controls (male n=6, female n=6). Individual-animal averages are indicated by circles for males and triangles for females. Error bars represent SEM around the estimated mean.

**Results of pairwise comparisons assessing outcome encoding on current trial (T0) given specified histories of reward over previous three trials (T -1, T-2, T -3).**

| | T0 Pairwise Comparison | T -1 Outcome | T -2 Outcome | T -3 Outcome | Z Ratio | p Value |
|---|---|---|---|---|---|---|
| mPFC-NAc | Rewarded vs Unrewarded | Unrewarded | Unrewarded | Unrewarded | 14.709144 | p<0.0001 |
| mPFC-NAc | Rewarded vs Unrewarded | Unrewarded | Unrewarded | Rewarded | 11.342379 | p<0.0001 |
| mPFC-NAc | Rewarded vs Unrewarded | Unrewarded | Rewarded | Unrewarded | 13.708781 | p<0.0001 |
| mPFC-NAc | Rewarded vs Unrewarded | Unrewarded | Rewarded | Rewarded | 12.042946 | p<0.0001 |
| mPFC-NAc | Rewarded vs Unrewarded | Rewarded | Unrewarded | Unrewarded | 9.963937 | p<0.0001 |
| mPFC-NAc | Rewarded vs Unrewarded | Rewarded | Unrewarded | Rewarded | 9.755649 | p<0.0001 |
| mPFC-NAc | Rewarded vs Unrewarded | Rewarded | Rewarded | Unrewarded | 6.794145 | p<0.0001 |
| mPFC-NAc | Rewarded vs Unrewarded | Rewarded | Rewarded | Rewarded | 5.635762 | p<0.0001 |
| vHip-NAc | Rewarded vs Unrewarded | Unrewarded | Unrewarded | Unrewarded | 9.6733623 | p<0.0001 |
| vHip-NAc | Rewarded vs Unrewarded | Unrewarded | Unrewarded | Rewarded | 5.9858316 | p<0.0001 |
| vHip-NAc | Rewarded vs Unrewarded | Unrewarded | Rewarded | Unrewarded | 5.4500847 | p<0.0001 |
| vHip-NAc | Rewarded vs Unrewarded | Unrewarded | Rewarded | Rewarded | 5.2443238 | p<0.0001 |
| vHip-NAc | Rewarded vs Unrewarded | Rewarded | Unrewarded | Unrewarded | 3.801793 | p=0.0011 |
| vHip-NAc | Rewarded vs Unrewarded | Rewarded | Unrewarded | Rewarded | 3.4495963 | p=0.0045 |
| vHip-NAc | Rewarded vs Unrewarded | Rewarded | Rewarded | Unrewarded | 1.2310677 | p=0.8605 |
| vHip-NAc | Rewarded vs Unrewarded | Rewarded | Rewarded | Rewarded | 0.8397556 | p=0.9834 |

Supplementary Table 1. Results of pairwise comparisons assessing outcome encoding on current trial (T0) given specified histories of reward over previous three trials (T -1, T-2, T -3).

**Linear mixed model of latency on subsequent trial based on mPFC-NAc and vHip-NAc activity at ITI end**

Model: Latency ~ mPFC * vHip + (1 | Animal ID) + (1 | Day), family = Gamma

| | Estimate | Standard Err( | t value | p value |
|---|---|---|---|---|
| Intercept | 0.459593 | 0.036952 | 12.438 | 2E-16 |
| PFC | -0.033814 | 0.004059 | -8.331 | 2E-16 |
| vHip | -0.030279 | 0.003969 | -7.629 | 2.36E-14 |
| PFC:vHip | 0.006913 | 0.00244 | 2.834 | 0.0046 |

Random Effects

| | Variance | Standard Deviation |
|---|---|---|
| Animal ID | 0.004705 | 0.06859 |
| Day | 0.006334 | 0.07959 |

Supplementary Table 30. Linear mixed model of latency on subsequent trial based on mPFC-NAc and vHip-NAc activity at ITI end. Latency ~ mPFC * vHip + (1IAnimal ID) + (1IDay), family = Gamma.
